# The influence of fluctuating population densities on evolutionary dynamics

**DOI:** 10.1101/444273

**Authors:** Hanja Pisa, Joachim Hermisson, Jitka Polechová

## Abstract

The causes and consequences of fluctuating population densities are an important topic in ecological literature. Yet, the effects of such fluctuations on maintenance of variation in spatially structured populations have received little analytic treatment. We analyze what happens when two habitats coupled by migration not only differ in their tradeoffs in selection but also in their demographic stability – and show that equilibrium allele frequencies can change significantly due to ecological feedback arising from locally fluctuating population sizes. When an ecological niche exhibits such fluctuations, these drive an asymmetry in the relative impact of gene flow, and therefore, the equilibrium frequency of the locally adapted type decreases. Our results extend the classic conditions on maintenance of diversity under selection and migration by including the effect of fluctuating population densities. We find simple analytic conditions in terms of the strength of selection, immigration, and the extent of fluctuations between generations in a continent-island model. While weak fluctuations hardly affect coexistence, strong recurrent fluctuations lead to extinction of the type better adapted to the fluctuating niche – even if the invader is locally maladapted. There is a disadvantage to specialization to an unstable habitat, as it makes the population vulnerable to swamping from more stable habitats.

## Introduction

Classic evolutionary theory behind maintenance of variation in natural populations focuses on the balance between selection, migration and mutation (Levene, 1953; Maynard Smith, 1970; Bulmer, 1972). Over decades of research, the conditions for maintenance of polymorphism have been analyzed for increasingly complex systems with many loci, alleles and demes (reviewed in Bürger 2014). Yet typically, population dynamics are considered fast enough so that they do not need to be modeled explicitly – an assumption that we relax in this work.

Natural populations are often out of equilibrium (Hastings, 2004). The discovery that large and irregular fluctuations can be driven by very simple dynamics (Lorenz, 1963) has led to a surge of interest in ecological theory. Both within- and between-species interactions, such as density dependence and predator-prey interactions can lead to large fluctuations in population sizes (May, 1972, 1974; Levins, 1979). Even chaotic dynamics, arising from overcompensating density dependence *via* intraspecific predation, have been successfully induced in an experimental population of flour beetles (Costantino et al., 1995). While, nonetheless, chaotic dynamics are believed to be rare (Thomas et al., 1980; Ellner and Turchin, 1995), fluctuations in population size are omnipresent. In natural populations, these can be caused by both intra- and inter-specific interactions, as well as purely exogenous abiotic factors such as weather (Framstad et al., 1997; Turchin et al., 2000; Coulson et al., 2001).

Oscillations in population size rarely affect the whole species uniformly. The size of the fluctuations may change through the species’ range and at any given time, some subpopulations can even go extinct. These local extinctions may affect the persistence of the whole population (Levins, 1969, 1970; Hanski and Ovaskainen, 2003). Yet, dispersal among subpopulations can strongly mitigate the extent of fluctuations, diminishing the risk of extinction in spatially structured populations (Den Boer, 1968; Reddingius and Den Boer, 1970; Roff, 1974). The whole (meta-)population becomes more stable as immigration from more populated patches pushes local population sizes away from zero (Hanski, 1985, 1991). Notably, even weak migration can dampen chaotic fluctuations arising due to overcompensating density dependence (Ruxton, 1994; Stone and Hart, 1999; Allen et al., 1993).

While dispersal stabilizes the ecological dynamics of a single species in the presence of disturbances, the effect of the interplay of dispersal and disturbances is more complex once we consider more species or within-species diversity (Holt, 1983b). Strong dispersal can lead to both global and local loss of variation, when some of the locally adapted types get swamped by other, more abundant variants. Interestingly though, diversity can increase with dispersal in the presence of disturbances and specific trade-offs in adaptation at low *vs*. high densities (*r*/*K*-selection, MacArthur and Wilson 1967; Pianka 1970). Even in the absence of spatial structure, coexistence on a single resource is possible if the trade-off in resource use leads to a stronger intra-specific competition than inter-specific competition (“relative nonlinearity”, Armstrong and McGehee 1980; Chesson 1994). When the balance between the ability to grow fast (high *r*) *vs*. efficiently (high *K*) can evolve, selection may favor intermediate values of both r and K (Roughgarden, 1971; Gadgil and Solbrig, 1972; Turelli and Petry, 1980; Lande et al., 2009; Engen et al., 2013; Lande et al., 2017). In contrast to well-mixed populations (Armstrong and McGehee, 1980; Turelli and Petry, 1980; Chesson, 1994), coexistence of multiple types on a single resource becomes considerably more robust when populations are spatially structured. Notably, in an environment with frequent local disturbances, coexistence of a quick (“weedy”) colonizer which grows fast from low densities and an efficient user of resources is possible under a broad range of conditions (Hastings, 1980; Comins and Noble, 1985; Tilman et al., 1994; Chesson, 2000; Long et al., 2007; Yodzis, 2013). In the presence of spatial structure, a wide range of *r*−*K* trade-offs readily leads to coexistence. In contrast to these previous studies, *r* and *K* are properties of the niches in our model: there is no trade-off in *r* and *K* which drives coexistence of multiple types when population densities fluctuate. We ask how coexistence between locally adapted types is influenced by out-of-equilibrium population dynamics.

We study coexistence of two discrete types (genotypes, species, bacterial clones) that are locally adapted to two habitats. These are coupled by migration and one (or both) of the habitats exhibit deterministic fluctuations in population size. When the environment varies across space, dispersal between two habitats brings in locally maladapted variants. How is local adaptation affected by the recurrent asymmetric gene flow arising in locally fluctuating populations? The balance between the trade-offs in selection, and between the flows among the independently-regulated habitats is of known importance: for stable polymorphism to be maintained, the stronger the migration between the niches, the tighter must be the symmetry between the selection underlying the negative trade-offs in fitness in the different niches (Maynard Smith, 1970; Bulmer, 1972; Bürger, 2014). Yet, to our knowledge, it is not known whether the classic predictions are robust when population dynamics within the niches are taken into account. We start by modeling evolution jointly with discrete-time ecological dynamics, where fluctuations arise from overcompensating density dependence under Ricker’s regulation (1954). We then proceed by analyzing a simplification with imposed fluctuations in population size. This leads to explicit analytical conditions which provide a qualitative insight into the effect of fluctuations in population densities on evolutionary dynamics – whether these are due to extrinsic disturbances or arise from complex ecological population dynamics.

## Models and methods

Population dynamics are modeled jointly with the dynamics of the change in frequency of two types adapted to two different habitats. Throughout, we interpret these types as two different alleles in a one-locus haploid system, although they could also represent two different species. The trade-off in selection is density-independent. A relatively complex model where local, endogenous density fluctuations arise due to overcompensating density dependence in discrete-time precedes a simplified version with exogenous fluctuations imposed on a subpopulation’s density. We start with numerical demonstrations of temporal dynamics, and progress to stability analyses in the presence of fluctuations. All results presented in the stability analyses are analytic – but lengthy and complicated expressions are evaluated numerically and visualized in graphics. The stability analysis was assessed *via* the leading eigenvalues of the corresponding systems, and by the conditions for a protected polymorphism, which – in the case of two haploids in two demes – implies global convergence (Karlin and Campbell, 1980). More details are provided in the supporting information (section S1).

### Density dependent regulation of population densities (endogenous fluctuations)

First, we address a two-niche model with migration and joint evolutionary and ecological dynamics, where the density-dependent population growth follows Ricker’s regulation (Ricker, 1954). In this model, a delayed feedback in density regulation can cause large fluctuations of the population size beyond the carrying capacity.

The life cycle starts with migration, where *m*_12_ denotes the forward migration rate from deme 1 into deme 2, i.e. *m*_12_ is the proportion of individuals that emigrate from deme 1. *m*_21_ is the forward migration rate in the opposite direction. Throughout our analyses, we will mainly focus on the complementary cases of symmetric migration, *m*_12_ = *m*_21_ = *m* ≤ 1/2, and unidirectional migration, *m*_12_ = 0 (Fig. 1).

**Figure 1:**
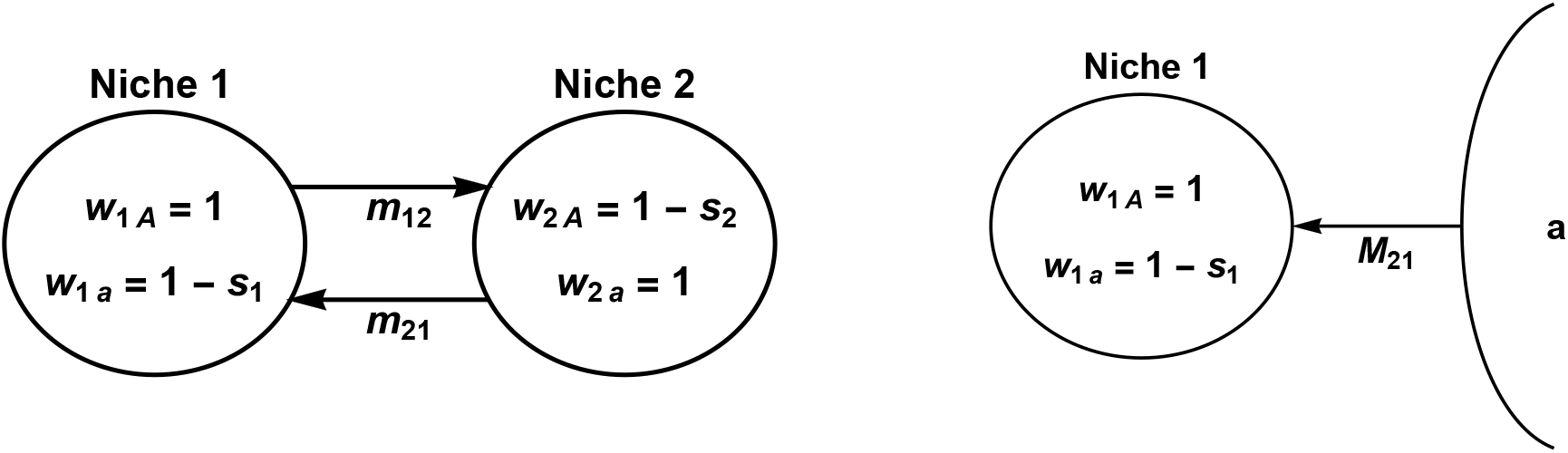
Evolution and maintenance of variation in the face of gene flow between populations in two divergent habitats. Left-hand side: Bidirectional migration between two ecological niches with forward migration rates *m*_12_ and *m*_21_. Type *A* is better adapted to niche 1; type *a* to niche 2. Right-hand side: Continent-island model with monomorphic immigration of the locally maladapted type *a* to the island. *M*_21_ denotes the absolute number of immigrants per generation.

At generation *t*, deme *i* is described by the population size, *N_i_*(*t*), and the frequency of the focal allele *A, p_i_*(*t*). We denote the second allele by *a*. For simplicity, primes label the intermediate variables after migration but before selection and population growth.

Following migration, the number of individuals in deme 1 is

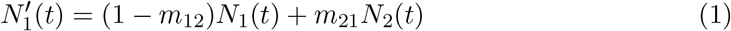

and the frequency of allele *A* in deme 1 is given by

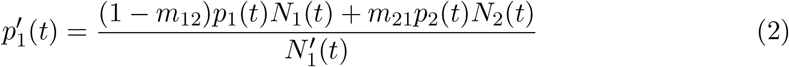

with similar recursions for deme 2.

There is a negative trade-off in the fitness of the alleles *A* and *a* in the two niches. The fitnesses of allele *A* are *w*_1*A*_ = 1 and *w*_2*A*_ = 1 − *s*_2_ in niches 1 and 2 respectively, and the corresponding fitnesses of allele *a* are *w*_1*a*_ = 1 − *s*_1_ and *w*_2*a*_ = 1 (see left-hand side of Fig. 1 for visualization). The selection coefficients *s*_1_, *s*_2_ range between 0 and 1. Thus, the focal allele *A* has a fitness advantage in niche 1 whereas *a* is better adapted to niche 2. In the next generation (after migration and selection), allele frequency of *A* in each deme *i* is

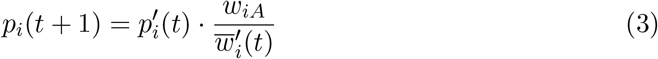

where 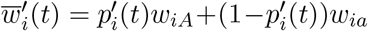 denotes the mean fitness after migration in deme *i*.

After migration, there is separate population regulation in each deme *i* following Ricker’s dynamics (Ricker, 1954):

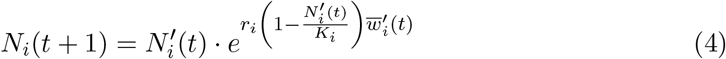

Here, *r_i_* denotes the intrinsic rate of increase, which gives the maximum growth rate of the population at low densities. Importantly, the rate *r_i_* is assumed to be a property of the niche rather than the types. This is a valid assumption when both types compete for the same resource within a niche. However, we suppose that the effective intrinsic rate of increase declines due to maladaptation (by multiplying the mean fitness 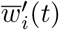 to the exponent): a maladapted population is not able to exploit the potential intrinsic rate of increase *r_i_*. The carrying capacity of a niche, though, remains unaltered by maladaptation and is given by *K_i_*. Throughout, we will consider niches with equal carrying capacity, i.e. *K* = *K*_1_ = *K*_2_, unless explicitly stated differently (notably, for the mainland-island model).

We are interested in the effects of fluctuations in population size around the carrying capacity on the evolutionary dynamics. In this study, the genetic composition of a population does not influence its carrying capacity (*soft selection*: Wallace 1975; Christiansen 1975), because this would add another layer of interactions between ecology and evolution to the system. We discuss an alternative model choice, where the mean fitness of a population affects its carrying capacity (*hard selection*) in the SI (section S7). The key results are the same for both models.

### Backward migration

For parts of our analysis it will prove helpful to rewrite the allele frequencies after migration to

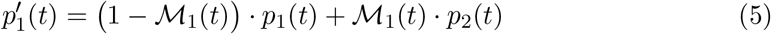

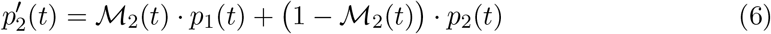

Here, 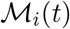 is the backward migration rate to deme *i*, which gives the proportion of new immigrants in the population of deme *i* at generation *t*. In our model, the backward migration rates are given by

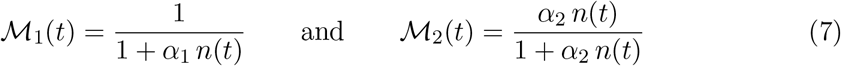

where *n*(*t*) = *N*_1_(*t*)/*N*_2_(*t*) is the proportion of the two population sizes, and *α*_1_ = (1 − *m*_12_)/*m*21 and *α*_2_ = *m*_12_/(1 − *m*_21_).

### Imposed population densities (exogenous fluctuations)

While fluctuations in the population size arising under high intrinsic rate of increase due to overcompensating density regulation such as over-exploitation of the resource or cannibalism are well documented (Symonides et al. 1986; Costantino et al. 1995; Turchin et al. 2000), it is useful to examine a simpler system where fluctuations in the population density are imposed.

In this second model, we impose fluctuations *D*_1_ in the population size of niche 1 such that it periodically fluctuates between two densities *K*_1_ − *D*_1_ and *K*_1_ + *D*_1_, whereas the population in niche 2 maintains a constant density *K*_2_ (Fig. S2). Again, we concentrate on the case *K* = *K*_1_ = *K*_2_. The equations for the evolutionary dynamics remain unchanged.

Finally, we analyze the most simplified setting with unidirectional migration from a monomorphic continent to an island which is subject to imposed fluctuations (see right-hand side of Fig. 1). Every generation, a constant absolute number *M*_21_ (corresponding to *m*_21_*K*_2_ = *m*_21_*K*_1_) of type *a* individuals migrate to the island, where they are locally maladapted (*w*_1*a*_ = 1 − *s*_1_). The population size on the island fluctuates between *K*_1_ + *D*_1_ and *K*_1_ − *D*_1_ in subsequent generations. (We assume that population regulation is strong enough to counteract the ecological effects of immigration.) For this model, we can find an explicit solution for the equilibrium frequency of the focal type *A* on the island (see SI, section S10) and compact analytical conditions for coexistence that allow for comprehensive interpretation.

## Results

We prove that in structured populations, the evolutionary dynamics strongly depend on whether the ecological dynamics maintain constant densities or exhibit fluctuations (which can arise from deterministic intrinsic dynamics or can be imposed extrinsically). As explained in detail in the methods, we study this phenomenon *via* a model of two discrete types living in two habitats with migration between them. There is a trade-off in adaptation to the two habitats and they also differ in their inherent productivity (which determines the attainable intrinsic rate of increase of the local population). Numerical demonstrations of the temporal dynamics – showing that allele frequencies can change significantly due to the ecological feedback – precede a stability analysis of the polymorphic equilibrium.

### Population dynamics affect evolution

When the density of a subpopulation fluctuates, allele frequencies evolve to a different equilibrium than predicted for population densities that are stable in time – see Fig. 2.

**Figure 2:**
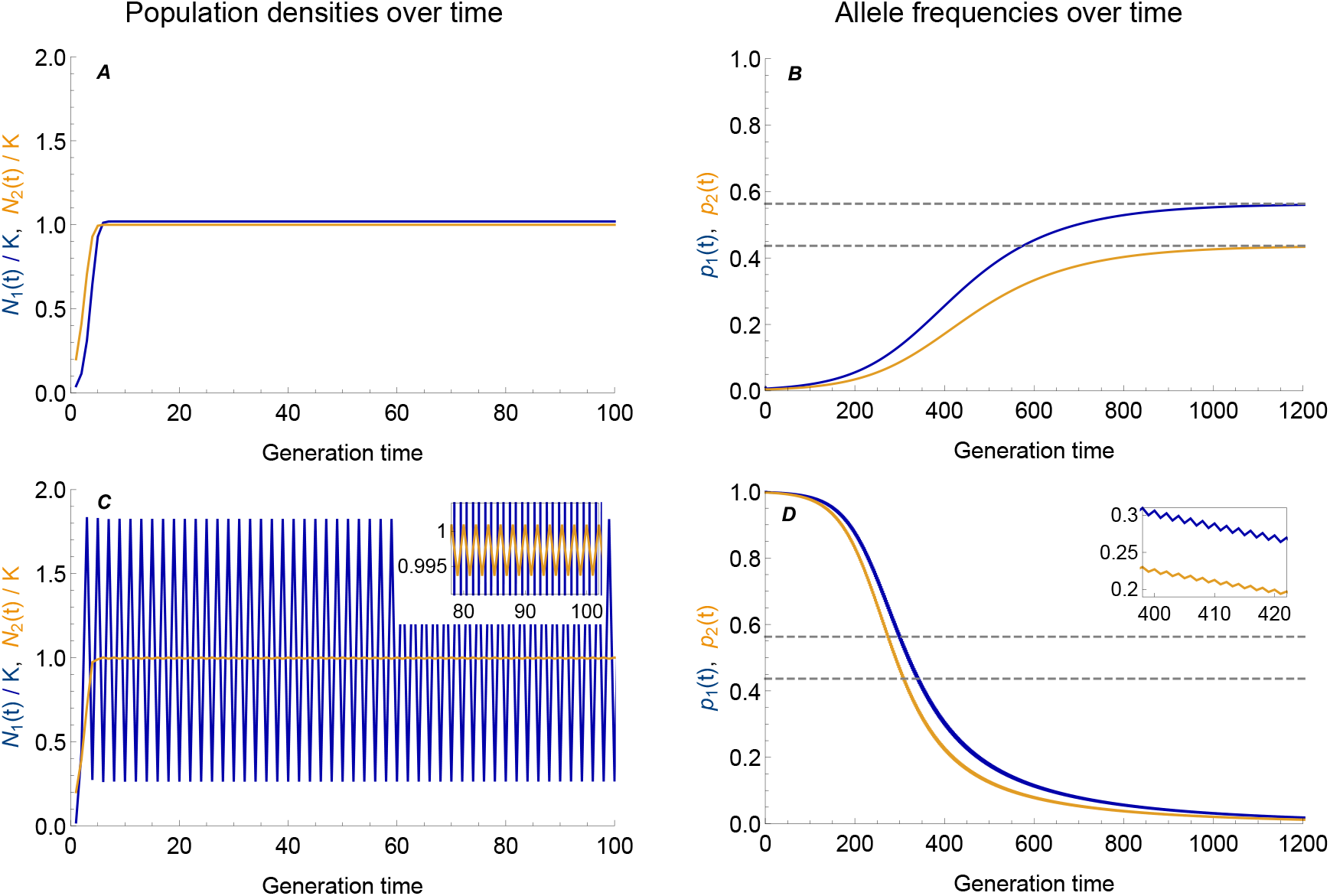
Ecological dynamics can change the evolutionary predictions. (A, C): Population dynamics in niche 1 (blue) and niche 2 (dark orange) following Ricker’s dynamics (Eq. 4) with low *vs*. high intrinsic rate of increase in niche 1 (A *r*_1_ = 1 *vs*. C *r*_1_ = 2.5). (B, D): The allele frequencies of the focal type, which is favored in niche 1, is shown in blue for niche 1 and in orange for niche 2. The dashed lines represent the expected equilibrium allele frequencies of the focal type in the absence of ecological dynamics. When population sizes evolve to a stable constant equilibrium (A), evolutionary dynamics behave as predicted (B): even when initially rare (*p*_1_(0) = 0.01, *p*_2_(0) = 0.005), the focal type is able to invade and its frequency converges to the globally asymptotically stable evolutionary equilibrium predicted in the absence of population dynamics (dashed lines). However, as the population density in niche 1 fluctuates due to overcompensating density dependence (C, blue), the frequency of the locally favored type decreases in both niches (D). Although initially very frequent (*p*_1_(0) = 0.999, *p*_2_ (0) = 0.997), the focal type eventually dies out due to the fluctuations in the niche where it is adapted to. The inset highlights the fluctuations in the allele frequencies, which are due to the fluctuating population density. In (A) the population sizes at equilibrium are equal (*N*_1_/*K* is offset by 0.02 for visual clarity). (C) In niche 2, intrinsic rate of increase is low enough (*r*_2_ = 1) so that the density stays nearly constant at carrying capacity (see inset) despite the strong fluctuations in niche 1. Parameters: *s*_1_ = *s*_2_ = 0.05, *m*_12_ = *m*_21_ = 0.1, *r*_1_ = 1 (A, B), *r*_1_ = 2.5 (C, D), *r*_2_ = 1, *K* = *K*_1_ = *K*_2_.

In the absence of ecological dynamics, we can predict the allele frequencies maintained at equilibrium (Eq. S4). These evolutionary predictions continue to be true when population dynamics are included as long as they lead to stable, constant densities (Fig. 2A, B). Whenever population densities tend to (and remain at) carrying capacity, allele frequencies converge to the unique, stable and globally attracting equilibrium predicted in the absence of ecology (Fig. 2B, dashed lines; Karlin and Campbell (1980) for the proof of global convergence). When the trade-off in selection is symmetric, this equilibrium is always polymorphic in the absence of fluctuations. Hence, even if the focal type is initially rare, it is able to invade and reach high frequencies.

When population dynamics do not lead to constant densities, the evolutionary predictions change. For high intrinsic rates of increase, overshooting of the carrying capacity is followed by overcompensating density regulation pushing the population to low densities, which leads to recurrent fluctuations in population size (Fig. 2C). The type adapted to a niche that exhibits such density fluctuations evolves to lower frequencies than predicted in the absence of ecology. Fig. 2D shows that a high intrinsic rate of increase in the first niche, which drives strong fluctuations in the local population size, is disadvantageous to the locally adapted type – and can ultimately result in its extinction. This is true even if the focal type is initially abundant and even if its equilibrium frequency in the absence of fluctuations is high (dashed lines). We analyze this effect in detail in the following paragraphs.

### How and why local fluctuations affect the evolutionary equilibrium

How does the equilibrium frequency of the focal type depend on the amplitude of fluctuations in population density? Fig. 3 shows the ecological (A) and evolutionary (B) dynamics at equilibrium for increasing values of the intrinsic rate of increase in niche 1, *r*_1_. For small *r*_1_, the population size reaches a stable equilibrium at carrying capacity (*N*_1_ = *K*_1_), and the allele frequencies converge to the stable migration-selection equilibrium predicted by classic population genetic theory. However, as *r*_1_ rises above a threshold, the ecological equilibrium becomes unstable and the population size in niche 1 fluctuates (blue lines in Fig. 3A). Simultaneously, the equilibrium allele frequencies of the type adapted to niche 1 start to decrease in both niches (blue, orange in Fig. 3B).

**Figure 3:**
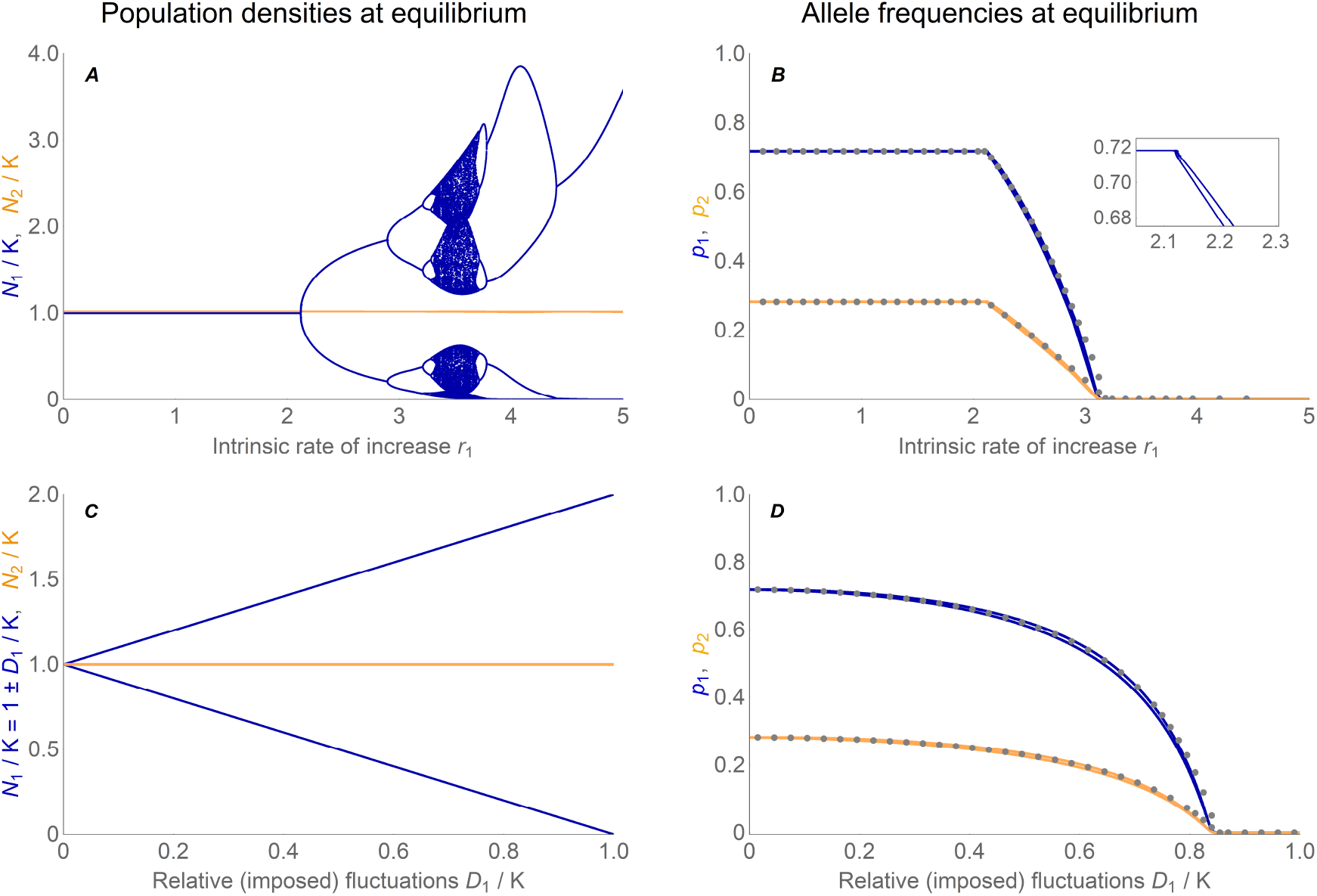
The evolutionary equilibrium changes continuously as fluctuations in population density increase. Fluctuations in local population size – whether they arise intrinsically due to overcompensating density dependence (A, blue) or are imposed extrinsically (C, blue) – lead to considerable changes in the evolutionary equilibrium of the focal type (B, D). The stronger the fluctuations in the density of one niche, the lower the equilibrium frequencies of the type that is adapted to this fluctuating habitat. (A) Fluctuations arise in niche 1 (blue) as the local intrinsic rate of increase, *r*_1_, increases above a threshold (see section S2). The population size in the second niche (orange) with a fixed intrinsic rate of increase, *r*_2_ = 1, stays close to the carrying capacity (*N*_2_/*K* is offset by 0.02 for visual clarity). The bifurcation diagram of the Ricker-regulated population (blue) differs from the classic one due to the stabilizing effect of dispersal (Fig. S3A). (B) As long as the intrinsic rate of increase *r*_1_ leads to stable population sizes, allele frequencies evolve towards the equilibrium predicted in the absence of joint population dynamics. However, with growing *r*_1_, fluctuations in the population density of niche 1 arise, and thus the equilibrium frequencies of the type favored in this niche decrease in both habitats (blue: *p*_1_ in niche 1, orange: *p*_2_ in niche 2). When fluctuations in the focal niche are strong, the locally adapted type goes extinct. Grey dots are an approximation of the evolutionary equilibrium, obtained by inserting the mean backward migration from the complete model (Fig. 4) into a simplified model with constant backward migration rates. This explains the evolutionary dynamics well unless fluctuations in backward migration rates are really large (see Fig. S6). The decrease in equilibrium frequency is smooth despite the complicated ecological dynamics, with only small fluctuations in the allele frequencies (inset). (C) This analogue to the branching diagram in (A) depicts the population density in niche 1 (blue) for linearly increasing values of imposed fluctuations (expressed relative to the carrying capacity as *D*_1_/*K*_1_). The population size in niche 2 stays constant (orange). (D) The equilibrium frequencies of the focal type decrease as a function of the (imposed) fluctuations in the niche where it is favored. Grey dots give the approximation using mean backward migration. The decrease in equilibrium frequencies is not as abrupt as it is in (B), because the increase in fluctuations is linear. Parameters: *s*_1_ = *s*_2_ = 0.1, *m*_12_ = *m*_21_ = 0.05, *K* = *K*_1_ = *K*_2_.

The larger the intrinsic rate of increase, the stronger are overshooting and overcompensation, which leads to oscillations of higher period and eventually to chaos, whenever migration is absent or really weak. In the absence of migration and selection, the transition to cycling occurs at *r*_1_ = 2 for the Ricker model (May and Oster, 1976). Yet, immigration from the second, stable deme stabilizes the ecological dynamics in two ways. First, by pushing the local population density away from zero: Even weak migration dampens fluctuations sufficiently so that the long-term dynamics no longer portrait the classic chaotic branching diagram of the single population Ricker model (Ruxton, 1994; Stone and Hart, 1999). Additionally, in our model, the effective intrinsic rate of increase 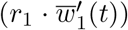 declines with the mean fitness due to immigration of maladapted types, increasing the threshold to cycling as a function of *r*_1_. A detailed analysis of the population dynamics, their stability analysis, branching points and how they are influenced by migration and selection can be found in the SI, section S2 (Fig. S3, S4).

Despite wildly fluctuating population dynamics with rising intrinsic rate of increase *r*_1_, the corresponding change in the polymorphic equilibrium is almost linear (Fig. 3A *vs*. B). The higher the variance in the population density of the focal niche, the lower the equilibrium frequency of the locally adapted type – until it dies out: when local fluctuations with increasing *r*_1_ become too strong, the stable polymorphism ceases to exist and the focal type goes extinct; the other type adapted to the stable niche fixes in both niches.

Also under a model with exogenous fluctuations in the focal niche (Fig. 3C), equilibrium frequencies of the locally adapted type decrease (Fig. 3D). Here, we assume that the external environment imposes periodic fluctuations between two densities (e.g. summer/winter) with constant average density *K*, but amplitude parametrized by *D*_1_. Fig. 3D shows a decline in the equilibrium frequencies of the locally adapted type with increasing amplitude of the fluctuations, which is entirely analogous to the effect of intrinsic fluctuations under Ricker’s regulation. In particular, the simplified model with imposed fluctuations demonstrates that the decrease in equilibrium frequency of the focal type is not simply caused by a drop in the mean population size within the focal niche when fluctuations arise.

What is the reason for the decrease in equilibrium frequency of the type that is locally adapted to the niche with oscillating population size? As both types grow reasonably well at low densities, the fluctuations in the focal niche can easily be exploited by types immigrating from the more stable environment, even if they are locally maladapted. In generations where the population density in the fluctuating niche is low, immigration brings in a significant number of individuals that are adapted to the other niche – resulting in a higher proportion of locally maladapted types in the focal niche. Afterwards, in the transition to a generation with high density, both types increase in number as the ecological dynamics are much faster than the evolutionary dynamics. In summary, in generations where the local density is low, the focal niche gets swamped by the other, more stable habitat. If fluctuations are strong, selection is not capable of compensating the recurrent swamping that occurs over a long time of repeated oscillations around the carrying capacity. Hence, the equilibrium allele frequency of the focal type decreases, i.e. the type that is adapted to the “better” niche, in which a high intrinsic rate of increase leads to fluctuations in density.

We can understand the increased swamping by gene flow due to fluctuations in local population densities in terms of backward migration rates. Backward migration measures the relative strength of immigration to a niche: it gives the proportion of individuals in the local population that immigrated from the other habitat. In contrast to forward migration, it depends on the population size in the focal niche. When densities fluctuate, they drive fluctuations in the backward migration rates (both to the fluctuating niche and to the other niche – light blue and light orange lines in Fig. 4): fluctuations up in local population size correspond to fluctuations down in the backward migration to this niche and *vice versa*. More importantly, fluctuations in the focal niche lead to an increase in its *average* backward migration rate (dark blue lines in Fig. 4). Indeed, whenever the population size in the focal niche is low, the backward migration to this niche is strongly increased due to the large contribution of immigrants relative to the locals. In contrast, the decrease in the backward migration rate in times of high local density is much smaller.

**Figure 4:**
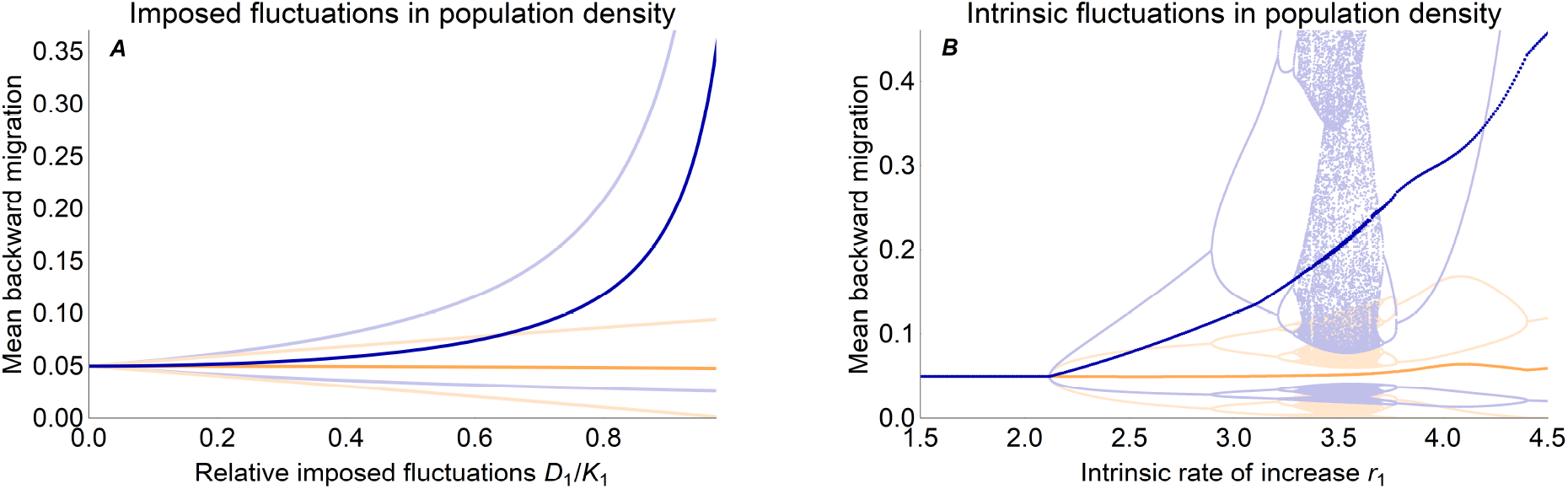
The mean backward migration to the focal niche increases significantly as fluctuations in the local population size grow. Because backward migration reflects immigration rates, it rises substantially when local population density is low. Thus, fluctuations in population size drive an increase in the average backward migration rate to the fluctuating niche (dark blue lines). (A) Under imposed density fluctuations in the focal niche, backward migration rates fluctuate up and down with a period of 2 (migration to niche 1 in light blue, niche 2 in light orange). As density fluctuations *D*_1_/*K*_1_ grow, the extent of these fluctuations in backward migration increases markedly for the focal niche (increasing mean, dark blue), but only about linearly for the stable niche 2 (barely declining mean, dark orange). (B) Under Ricker’s regulation, complex fluctuations in backward migration rates arise (light lines) as a high intrinsic rate of increase *r*_1_ drives fluctuations in population density. Again, this leads to a strong increase in mean backward migration to the focal niche, while hardly affecting the one to the stable niche (dark blue *vs*. dark orange). Dark lines were obtained by averaging backward migration over generations 1000 – 1300. The bump in the mean backward migration rates around *r*_1_ = 4 comes from the population size fluctuating to high densities in this regime. Parameters of selection and migration are the same as in Fig. 3.

The rise in “average relative immigration” as density fluctuations grow explains most of the change in the evolutionary equilibrium: using the mean backward migration rates as an approximation for fluctuations leads to a good estimate of the equilibrium allele frequencies under fluctuating population densities (*c.f*. grey dots in Fig. 3). We obtain this close approximation by replacing the migration rates in a model without joint population dynamics by the mean backward migration from the model with joint ecological dynamics (and density fluctuations). Hence, the mean backward migration gives a good measure for the strength of fluctuations and their impact on the evolutionary dynamics – especially when fluctuations in backward migration rates are not too strong (SI, section S3.2). When fluctuations in backward migration increase with intensifying density fluctuations, their small effect on equilibrium allele frequencies becomes apparent – and the approximation deviates slightly from the complete model (just before the focal type dies out in Fig. 3).

Stronger selection in a fluctuating niche counteracts higher mean backward migration rate. It makes the locally adapted type less vulnerable to the recurrent swamping, because the type’s ability to compete in the growing phase increases with selection. This holds true even when the maladapted type dominates in frequency after migration and even if the trade-off in selection between the niches is symmetric (i.e. also strong in the stable niche). Strong symmetric selection, albeit leading to less variation in both niches, stabilizes the polymorphism and maintains diversity for higher intrinsic rate of increase (Fig. S7).

The decrease in allele frequency of the type which is best adapted to a fluctuating environment is due to the combination of the ecological instability and immigration of a type adapted to an other, more stable habitat. In the absence of gene flow, the locally adapted type converges to fixation regardless of fluctuations. Given a constant immigration rate, increasing emigration may be essential for survival of the type adapted to the fluctuating niche (see Fig. S8). As fluctuations grow, higher asymmetry between emigration and immigration is necessary for the focal type to persist – the other niche acts as a reservoir of the focal type although it is locally maladapted. Increasing symmetric migration is only advantageous to the focal type under specific conditions, such as when the focal niche is larger (Fig. S9).

### Robustness to further scenarios

So far, we have assumed that one of the niches undergoes fluctuations in population density while the second niche is stable, but our findings generalize to oscillations in both niches. When both niches exhibit density fluctuations, the type which is adapted to the more stable habitat (where fluctuations are the smallest), increases in frequency (see Fig. 5). Specifically, when the population density in the second niche fluctuates, the focal type *A* benefits from these fluctuations as long as the first niche maintains a more stable density (*r*_1_ < *r*_2_). This relative advantage of the focal type declines, and so does its equilibrium frequency, as the amplitude of the fluctuations in the focal niche rises. When the intrinsic rates of increase are the same in both niches and population sizes are fluctuating but not yet chaotic, allele frequencies converge to about the same value as in the absence of fluctuations (intersection of gray dashed lines in Fig. 5B, D, F). This is because mean backward migration rates are identical in both niches when density fluctuations are the same. Similarly, the less fluctuating niche is subject to the weakest average backward migration, and therefore the type adapted to it swamps niches that undergo stronger fluctuations. This generalizes the phenomenon described in the previous paragraphs: when one niche is significantly less stable than the other niche, the locally adapted type goes extinct.

**Figure 5:**
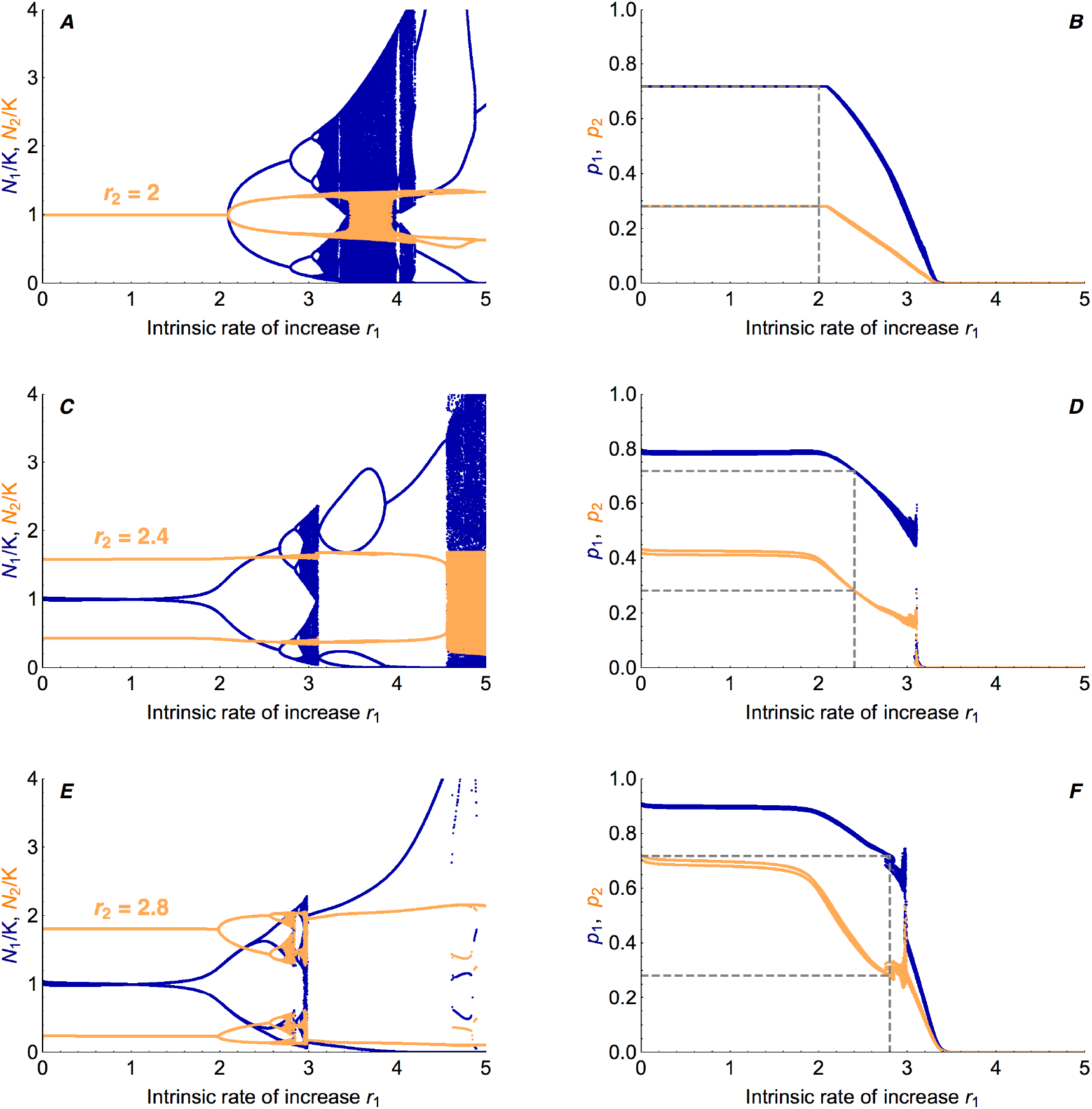
When both niches exhibit fluctuations, the type better adapted to the more stable niche increases in frequency. With growing intrinsic rate of increase in niche 2 (from top to bottom: *r*_2_ = 2, 2.4, 2.8), fluctuations in population density in niche 2 arise and intensify (A, C, E). This leads to higher equilibrium frequencies of the focal type (B, D, F) because it is able to swamp niche 2 whenever the population density there is low. Blue lines show the long-term behavior in niche 1, orange lines describe niche 2. Gray dashed lines indicate the diversity maintained if population densities were constant (horizontal), and the value of *r*_2_ (vertical). When the population density in niche 1 stays close to the carrying capacity while niche 2 exhibits fluctuations (*r*_2_ > 2, *r*_1_ < *r*_2_ – note that the branching point in niche 1 depends on the amplitude of fluctuations in niche 2), the focal type reaches higher equilibrium frequencies than predicted in the absence of fluctuations (horizontal dashed lines in D, F). The relative advantage of the focal type vanishes once *r*_1_ ≈ *r*_2_ (vertical dashed lines in D, F), where equilibrium frequencies are about the same as for constant population densities (intersection of vertical with horizontal dashed lines in B, D, F). When *r*_1_ > *r*_2_ (right of vertical dashed lines), niche 1 is less stable than niche 2, which leads to a decrease in the equilibrium frequency of the focal type – and eventually, as *r*_1_ increases, to its extinction. Parameters: *s*_1_ = *s*_2_ = 0.1, *m* = 0.05

The decline in equilibrium frequency of the type adapted to the less stable niche is independent of the underlying reason for the occurrence of fluctuations. For the evolutionary dynamics, only net fluctuations in population size matter – whether imposed or driven by intrinsic dynamics. Thus, the change in equilibrium frequency is robust also under other forms of population dynamics such as logistic density dependence (see SI: section S6, Fig. S10, S11). Here, fluctuations are weaker than under Ricker’s regulation, and we see that the focal type’s equilibrium frequencies decrease continuously even if polymorphism can be maintained under chaotic densities.

Our results are also robust to a model of hard selection, where the mean fitness of a population impacts both its growth rate and its carrying capacity. We discuss this alternative formalization in detail in the SI (section S7): the decline in equilibrium allele frequency with growing fluctuations is even steeper, because mean fitness in the fluctuating niche decreases due to a higher proportion of maladapted immigrants (Fig. S12).

Furthermore, the observed evolutionary phenomenon is qualitatively independent of whether migration precedes selection and population growth (as modeled here) or succeeds them (Fig. S13).

### Conditions for stable polymorphism

How do the classic conditions for maintenance of diversity under migration and selection change when local population size fluctuates? As long as population dynamics maintain constant densities, simple explicit conditions for a stable polymorphism can be derived (SI, section S1.1). For the general case of arbitrary migration rates and potentially unequal niche sizes these can be found in the SI (Eqs. S5, S6). The condition for coexistence of the two types when population densities are stable (and equal) and migration is symmetric, is given by

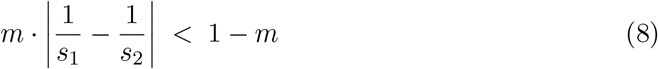

We see that both symmetry in selection strength between the niches and low migration stabilize the polymorphism. This balance between migration and selection expresses a necessary and sufficient condition for maintenance of polymorphism in the case of two haploids inhabiting two demes in the absence of population dynamics (or, when they are included but lead to a stable, constant ecological equilibrium). The formula is similar to the well-known sufficient conditions for maintenance of polymorphism in diploids and models in continuous time (Maynard Smith, 1970; Bulmer, 1972; Lenormand, 2002; Bürger, 2014). Condition (8) is visualized by the dashed lines in Fig. 6 (also Fig. S1): the stronger the asymmetry in selection, the larger selection needs to be relative to migration for diversity to be maintained. In the absence of fluctuations, if selection is strongly asymmetric, and weak relative to migration, there is no polymorphic equilibrium and allele frequencies converge to fixation.

**Figure 6:**
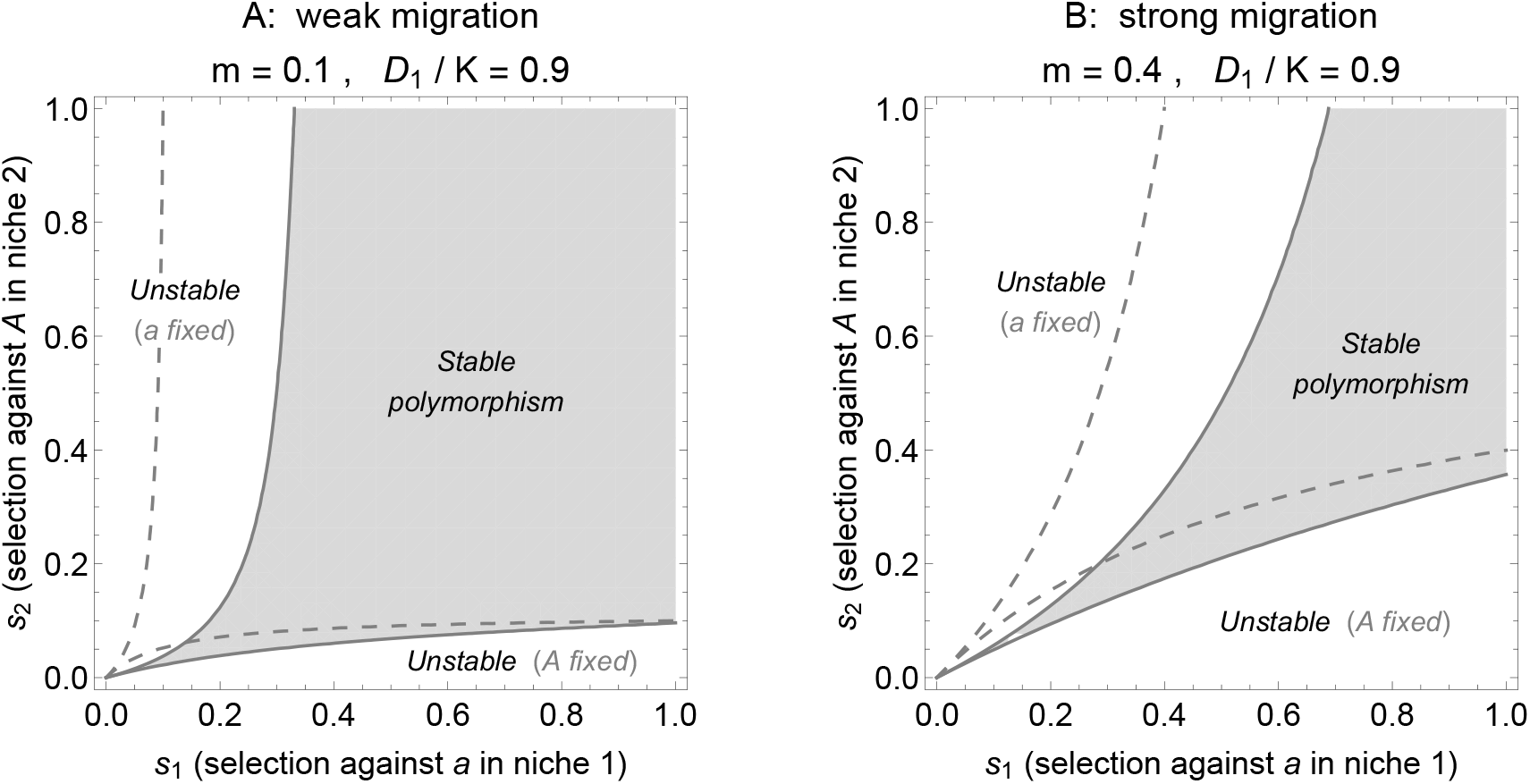
Fluctuations in the density of niche 1 shift and skew the parameter region where variation is maintained. Dashed lines give the condition for maintenance of polymorphism in the absence of fluctuations (*m* · |1/*s*_1_ − 1/*s*_2_| < 1 − *m*), while solid lines delimit the region where polymorphism is stable (grey area) under strong imposed fluctuations in the population density in niche 1 (*D*_1_/*K* = 0.9). To prevent extinction of the type adapted to the fluctuating niche, its selective advantage must be comparatively higher than when the densities are stable: the stability region shifts to the right, at a rate which is nearly independent of the strength of migration (A *vs*. B). Fluctuations broaden the parameter space that leads to fixation of the type *a* adapted to the stable niche by increasing the pressure on the focal type *A*. However, if asymmetry in selection favors the focal type (*s*_1_ ≫ *s*_2_), then fluctuations counteract that asymmetry and polymorphism is maintained more easily: the region where type *A* gets fixed shrinks.

When population densities fluctuate, the explicit analytic conditions for a globally stable polymorphism are long and complex (SI, section S1.3). They are derived using stability analysis of the model with imposed fluctuations in niche 1 – where this analysis is possible although we do not have an explicit formula for the equilibrium allele frequencies under fluctuating population densities. The conditions for maintenance of diversity under strong imposed fluctuations in the focal niche are shown in Fig. 6 (solid lines, encasing the grey area).

The parameter region where a stable polymorphic equilibrium is maintained changes when a subpopulation’s density fluctuates (Fig. 6: solid *vs*. dashed lines) – but the stabilizing effect of weak migration and strong selection is preserved under both constant and fluctuating population dynamics. Variation is maintained more easily when dispersal is low (Fig. 6A *vs*. B), and the robustness of the polymorphism broadens as symmetric selection gets stronger (polymorphism is maintained for larger fluctuations as selection intensifies). Therefore, the shape of the stability region is fairly robust to fluctuations in density. However, fluctuations in one niche, while the other niche is stable, introduce an asymmetry in the range of selection coefficients for which variation is maintained. The stability region is shifted because selection against the focal type and density fluctuations in the focal niche have a similar impact on the evolutionary dynamics: both lead to a decrease in equilibrium allele frequency of the focal type.

Under fluctuations in the focal niche, more variation is maintained when the selection trade-off between the niches is asymmetric and benefits the focal type (*s*_2_ < *s*_1_). Fluctuations in the first niche lower the equilibrium frequency of the locally favored type (*A*), while increasing the proportion of the other type (*a*). Therefore, they reduce the parameter region where the focal type *A* becomes fixed (Fig. 6: white area, bottom right). This means, whenever the focal type *A* has a strong relative selective advantage (*s*_2_ ≪ *s*_1_), fluctuations in the niche where it is favored counteract the asymmetry in selection by increasing the proportion of the disadvantaged type *a*. In that regime, this broadens the region where polymorphism is maintained: density fluctuations can support diversity if asymmetry in selection trade-offs favors the type adapted to the fluctuating habitat. On the other hand, when asymmetry in selection trade-offs imposes a strong disadvantage on the focal type *A* (*s*_2_ ≫ *s*_1_), its allele frequencies decrease and fluctuations lead easily to its extinction. Then fluctuations intensify the asymmetry in selection. Hence, a significantly larger range of parameters leads to fixation of the other type, *a*, when fluctuations occur in the first niche (Fig. 6: white area, top left). In summary, in the presence of fluctuations, polymorphism is maintained for stronger selection against maladapted immigrants (*s*_1_), and weaker selection against the focal type in the stable niche (*s*_2_).

### Polymorphism under imposed fluctuations in a continent-island model

Since the analytic conditions for maintenance of polymorphism under imposed fluctuations are extensive and complicated in the model with bidirectional migration (visualization in Fig. 6), it proves useful to analyze a continent-island model with imposed fluctuations on the island. In the SI we demonstrate that the effect of (intrinsic) density fluctuations on the evolutionary dynamics is robust to whether migration is symmetric or unidirectional to the fluctuating niche: the equilibrium frequency of the focal type is simply shifted to lower values under monomorphic immigration (section S9). For the continent-island model, simple explicit conditions for the persistence of the resident type on the island can be derived.

We obtain conditions for persistence of the locally adapted type in terms of selective disadvantage of the immigrating type, *s*_1_, the relative fluctuation size on the island, *D*_1_/*K*_1_, and the “rate of immigration” – the mean ratio of immigrants to locals (at the beginning of the life cycle), *M*_21_/*K*_1_ = *m*_21_.

Extinction of the locally adapted resident type on the island is certain whenever the selective disadvantage of the immigrating type is not strong enough to prevent swamping by the continent. Namely, for survival of the resident type to be possible, the relative strength of immigration at the time of selection, *M*_21_/(*K*_1_ + *M*_21_) = *m*_21_/(1 + *m*_21_), needs to be weaker than the local selective disadvantage of the immigrants, *s*_1_:

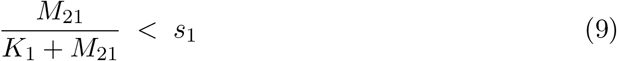

This condition recovers the known threshold for survival of a locally adapted type in a continent-island model in the absence of population dynamics (Haldane, 1930; Wright, 1931).

In the presence of fluctuations, condition (9) is necessary but not sufficient. The resident type persists on the island if and only if

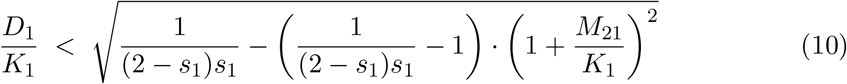

This threshold, which we denote by (*D*_1_/*K*_1_)*, gives the critical relative fluctuation size at which the resident type dies out. It increases with the selection coefficient *s*_1_ and decreases with the rate of immigration *M*_21_/*K*_1_, as illustrated in Fig. 7A.

**Figure 7:**
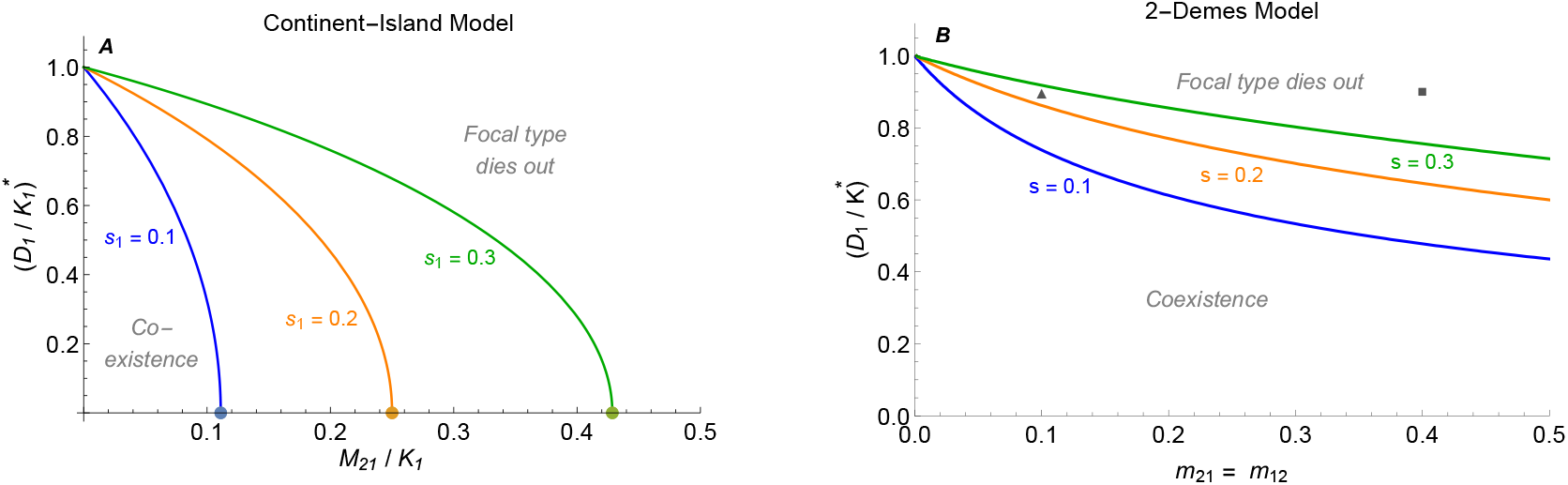
Strong fluctuations lead to extinction even when migration is weak, but the effect is attenuated under bidirectional migration. The critical relative fluctuation size is a decreasing function of the migration rate and an increasing function of selection (*s*_1_ = 0.1 (blue), 0.2 (orange), 0.3 (green)). (A) In the continent-island model, locally maladapted immigrants swamp the island when immigration is too strong. Just once the relative immigration falls below the critical value (dots: *M*_21_/*K*_1_ = *s*_1_/(1 − *s*_1_), see Eq. 9), coexistence becomes possible. Then the critical relative fluctuation size (*D*_1_/*K*_1_)* increases abruptly: weak fluctuations have a weak effect on the maintenance of diversity on the island. When migration is weak relative to selection (*M*_21_/*K*_1_ ≪ *s*_1_/(1 − *s*_1_)), the dependency between critical relative fluctuation size and the rate of immigration becomes almost linear – the stronger the selection against the immigrating type, the more is the resident type robust to fluctuations. (B) In the 2-demes model, migration can be polymorphic, and hence even strong dispersal does not necessarily lead to extinction of the focal type. The lines depict the critical relative fluctuation size when migration and selection are symmetric (*m*_12_ = *m*_21_, *s*_1_ = *s*_2_). The triangle (*m* = 0.1) and square (*m* = 0.4) provide a reference to Fig. 6 which depicts the stability of the system for variable selection (*s*_1_ ≠ *s*_2_) but a fixed value of fluctuations (*D*_1_/*K* = 0.9).

Condition (10) implies and extends the threshold for swamping in the absence of population dynamics (Eq. 9). The critical relative fluctuation size (*D*_1_/*K*_1_)* is a positive real number, whenever condition (9), which is equivalent to *M*_21_/*K*_1_ < *s*_1_/(1 − *s*_1_), is fulfilled. The critical relative fluctuation size equals zero when *M*_21_/*K*_1_ = *s*_1_/(1 − *s*_1_), and increases to positive values as the rate of immigration decreases below this threshold for swamping. In summary, whenever the relative number of immigrants is small enough (Eq. 9), coexistence becomes possible, but depends on the amplitude of fluctuations on the island (Eq. 10).

This result corresponds to our previous findings: the fewer locally maladapted immigrants arrive on the island and the stronger their maladaptation, the higher the frequency of the resident type in the absence of fluctuations. Therefore, the locally adapted type becomes more robust to fluctuations. In general, small fluctuations do not yet lead to extinction of the resident type once the balance between selection and immigration allows for coexistence of the two types (see Fig. 7A). Weak fluctuations have a weak effect on the maintenance of diversity on the island.

There are two cases when fluctuations can be extensive without leading to extinction of the resident type: when immigration tends to zero, or when selection against the immigrants is lethal, *s*_1_ → 1. These are the scenarios where the island is monomorphic for the resident type because the immigrants cannot survive there. As immigration increases from zero, the critical relative fluctuation size decreases almost linearly, with a negative rate of −1/((2 − *s*_1_)*s*_1_) + 1 (Fig. 7A and Fig. S15). This rate flattens as selection against the immigrating type increases. Thus, the critical relative fluctuation size declines faster with increasing immigration when selection is weak. Conversely, the stronger the selection, the larger are the fluctuations under which the resident type is still able to survive – a phenomenon that we discussed in the model with symmetric migration and density dependent population regulation already (Fig. S7).

How does the critical relative fluctuation size in the continent-island model compare to the critical relative fluctuation size in the model with two niches and symmetric migration between them (Fig. 7B)? When there is bidirectional migration between two niches, fluctuations in the focal niche are less detrimental to the locally adapted type than in the continent-island model (Fig. 7A *vs*. B). Then, the other more stable niche is inhabited by both types, leading to a polymorphic (rather than monomorphic) immigration into the focal niche. Hence, migration out of the fluctuating niche can be essential for persistence of the locally adapted type: the ability to find refuge in a stable niche is strongly beneficial for its maintenance – even if it is maladapted to that other niche and would go extinct there in the absence of immigration.

## Discussion

In nature, the productivity and stability of environments vary through space. Species live across sets of heterogeneous habitats that differ both in their demographic stability and evolutionary optima. For example, many species live both in their natural habitat as well as in strongly human-altered environments such as parks, fields or houses. These habitats favor different adaptation, and typically differ in their stability as well. Likewise, peripheral populations of species often experience more extreme and unstable environments, inducing fluctuations in population sizes. Our results demonstrate that in structured populations, recurrent density fluctuations within an ecological niche have a substantial effect on the diversity of the whole population. Whenever local population size is low, the locally adapted type is vulnerable to swamping by migration from a neighboring habitat. The asymmetry in gene flow induced by fluctuating populations is reflected by an increase in average backward migration rate, which measures the mean relative immigration into the focal niche. We obtain simple explicit conditions for persistence of the resident type and show that while small fluctuations hardly affect coexistence, even weak migration may drive the locally adapted type to extinction if fluctuations are strong. Stability in population size can be just as important for maintenance of diversity as the selective advantage within a niche.

In the 60s and 70s, ecological instability and chaotic dynamics were in the focus of attention (May, 1972, 1974; Levins, 1979), and the fact that migration can balance local disturbances has been recognized early on: “The effect of extreme conditions in one place will be leveled out to some degree by less extreme conditions in others. Migration can contribute to the leveling influence of spatial heterogeneity.” (Den Boer, 1968). Later, Chesson (1985) formalized these ideas of coexistence as a “spatial storage effect” – coexistence is facilitated because differing micro-habitats buffer a species against poor recruitments that occur during periods when the other species has a competitive advantage. In our work, local population sizes fluctuate, but within each habitat, evolutionary optima are stable across time. While dispersal stabilizes the population dynamics, it has a destabilizing effect on the maintenance of polymorphism: weak migration across a heterogeneous habitat increases local variation, but polymorphism cannot be maintained when immigration is too high (reviewed in Lenormand 2002). When population densities fluctuate, swamping by migration becomes powerful, and the equilibrium frequency of the type which is adapted to the “fluctuating” niche decreases. There is a disadvantage to adaptation to an ecological niche which exhibits fluctuations in population size.

Migration between habitats stabilizes population dynamics – and can even suppress chaotic fluctuations (Allen et al., 1993; Stone and Hart, 1999) – for types which share a common resource. Yet, due to ongoing migration, types adapted to unstable, strongly fluctuating niches will be replaced by their competitors, which benefit from migration from a more stable environment. The replacement itself may have a minor stabilizing effect on the ecological dynamics when the effective growth rate decreases with maladaptation (Fig. S3B). As long as the habitats are connected by dispersal, fluctuations of the persisting type remain suppressed – even if in the absence of population structure, these would be chaotic.

It is an important assumption of our model that the intrinsic rate of increase is a property of the niche rather than the genotype – as is the case when multiple types use a common abundant resource. Similarly, exogenous fluctuations are imposed on the entire population within a niche. As the fluctuations of a niche apply to its whole subpopulation, they lead to an increase in relative immigration which is independent of evolution and applies to all genotypes at the same rate. Therefore, the recurrent swamping overcomes the evolutionary trade-off and the type adapted to the most severely fluctuating niche gets swamped by migration from more stable habitats. Its allele frequency declines slowly over many generations of fluctuations in population density. In ecological models where the capacity to grow well from low densities and the trade-off between the types are combined to one parameter (Smith, 1998; Luís et al., 2011), we do not observe the effect that fluctuations in the resource-rich niche ultimately harm the locally more competitive type. This is because density fluctuations then only arise due to a dual advantage of this type. The coupling of demography and evolution is an essential feature of our model.

Here, we assume soft selection, so that the carrying capacity does not change with maladaptation. Under hard selection, both intrinsic rate of increase and carrying capacity change with the mean fitness of the population. Then, the decline in frequency of the type adapted to the unstable environment is slightly faster with growing fluctuations (see Fig. S12), because the asymmetry in average gene flow increases as the carrying capacity in the focal niche decreases. We have started with the Ricker model of a population’s density dependent growth, as it is established in theoretical literature and there are examples where it gives good fit to experimental data (Thomas et al., 1980). Here, fluctuations driven by delayed feedback in density regulation arise from discrete time dynamics. However, our main results are robust to extrinsically imposed fluctuations and alternative choices of density regulation such as logistic density dependence (see Fig. 3 and Fig. S11). To a good approximation, the effect of fluctuations can be recovered using mean backward migration rates. Therefore, our results obtained from discrete-time dynamics should hold for continuous time as well – as long as the differences in the time scales of fluctuations in the ecological *vs*. evolutionary dynamics are similar as in our model.

Furthermore, although we restrict ourselves to the study of two types inhabiting two heterogeneous niches, we are confident that our results generalize to *n* types in *m* niches with various trade-offs in fitness between the niches. The deleterious effect of fluctuations would then be observable as a decrease in equilibrium frequency of the type(s) that is/are best adapted to the environment(s) that exhibit(s) fluctuations in population density. We expect a decrease in equilibrium frequency of the type adapted to the niche exhibiting the most severe fluctuations. However, when one type is adapted to multiple unstable habitats, migration between them can stabilize the population dynamics (Allen et al., 1993; Ruxton, 1994; Stone and Hart, 1999) and therefore increase the type’s robustness against gene flow from neighboring habitats with trade-offs in adaptation.

Our results are of particular relevance for maintenance of local adaptation to peripheral populations (Holt, 1983a,b; Lennon et al., 1997; Sexton et al., 2009; Holt and Barfield, 2011), which often experience more severe fluctuations than central habitats (Harrison, 1991). Our deterministic model predicts that if genetic drift is weak relative to selection, adaptation to local conditions is considerably more difficult when population density fluctuates strongly. Then, dispersal can even prevent local adaptation. However, our predictions could change when we include stochasticity: by keeping the locally adapted type away from very low densities, migration could reduce stochastic extinction risk, as even weak dispersal has a strong stabilizing effect on density-dependent population dynamics. Therefore, when genetic drift is strong, dispersal could be beneficial even when it brings in also maladapted types. A similar effect is seen in models of evolution to species range margins with a continuously varying resource – once genetic drift becomes important, increasing migration rate improves the adaptation to marginal conditions (Polechová, 2018). The effects of genetic and demographic stochasticity in our simple model, as well as the impact of added instability in peripheral populations on adaptation to marginal conditions, would be an interesting subject to dedicated future study.

Understanding the impact of fluctuations is becoming progressively more important to conservation biology. Extreme weather, which currently appears increasingly often as the stability of the climate is decreasing, can readily induce larger fluctuations in population densities. We show that even when population sizes stay large enough so that stochastic extinction is not yet a concern, the variance in population size is an important factor for maintenance of diversity. As an omnipresent phenomenon in natural populations, the effects of fluctuations on diversity has also come into focus of experimental studies (Buckling et al., 2000; Rainey et al., 2000; Buckling et al., 2007). With our work, we aim to elucidate the effects of unstable population dynamics, and highlight the importance of considering ecology and evolution jointly.

## Supporting information

Supplementary text including all figures

## References

Allen, J., W. Schaffer, and D. Rosko, 1993. Chaos reduces species extinction by amplifying local population noise. Nature 364:229.

Armstrong, R. A. and R. McGehee, 1980. Competitive exclusion. The American Naturalist 115:151–170.

Buckling, A., M. Brockhurst, M. Travisano, and P. Rainey, 2007. Experimental adaptation to high and low quality environments under different scales of temporal variation. Journal of Evolutionary Biology 20:296–300.

Buckling, A., R. Kassen, G. Bell, and P. Rainey, 2000. Disturbance and diversity in experimental microcosms. Nature 408:961–964.

Bulmer, M., 1972. Multiple niche polymorphism. The American Naturalist 106:254–257.

Bürger, R., 2014. A survey of migration-selection models in population genetics. Discrete & Continuous Dynamical Systems-B 19:883–959.

Chesson, P., 1994. Multispecies competition in variable environments. Theoretical Population Biology 45:227–276.

Chesson, P., 2000. General theory of competitive coexistence in spatially-varying environments. Theoretical population biology 58:211–237.

Chesson, P. L., 1985. Coexistence of competitors in spatially and temporally varying environments: a look at the combined effects of different sorts of variability. Theoretical Population Biology 28:263–287.

Christiansen, F. B., 1975. Hard and soft selection in a subdivided population. American Naturalist 109:11–16.

Comins, H. and I. Noble, 1985. Dispersal, variability, and transient niches: species coexistence in a uniformly variable environment. The American Naturalist 126:706–723.

Costantino, R. F., J. M. Cushing, B. Dennis, and R. A. Desharnais, 1995. Experimentally induced transitions in the dynamic behaviour of insect populations. Nature 375:227.

Coulson, T., E. A. Catchpole, S. D. Albon, B. J. Morgan, J. Pemberton, T. H. Clutton-Brock, M. Crawley, and B. Grenfell, 2001. Age, sex, density, winter weather, and population crashes in soay sheep. Science 292:1528–1531.

Den Boer, P. J., 1968. Spreading of risk and stabilization of animal numbers. Acta biotheoretica 18:165–194.

Ellner, S. and P. Turchin, 1995. Chaos in a noisy world: new methods and evidence from time-series analysis. The American Naturalist 145:343–375.

Engen, S., R. Lande, and B.-E. Sæther, 2013. A quantitative genetic model of r-and k-selection in a fluctuating population. The American Naturalist 181:725–736.

Framstad, E., N. C. Stenseth, O. N. Bjørnstad, and W. Falck, 1997. Limit cycles in norwegian lemmings: tensions between phase–dependence and density-dependence. Proceedings of the Royal Society of London B: Biological Sciences 264:31–38.

Gadgil, M. and O. T. Solbrig, 1972. The concept of r-and k-selection: evidence from wild flowers and some theoretical considerations. The American Naturalist 106:14–31.

Haldane, J. B. S., 1930. A mathematical theory of natural and artificial selection. (part vi, isolation.). Mathematical Proceedings of the Cambridge Philosophical Society 26:220–230.

Hanski, I., 1985. Single-species spatial dynamics may contribute to long-term rarity and commonness. Ecology 66:335–343.

Hanski, I., 1991. Single-species metapopulation dynamics: concepts, models and observations. Biological Journal of the Linnean Society 42:17–38.

Hanski, I. and O. Ovaskainen, 2003. Metapopulation theory for fragmented landscapes. Theoretical population biology 64:119–127.

Harrison, S., 1991. Local extinction in a metapopulation context: an empirical evaluation. Biological journal of the Linnean Society 42:73–88.

Hastings, A., 1980. Disturbance, coexistence, history, and competition for space. Theoretical Population Biology 18:363–373.

Hastings, A., 2004. Transients: the key to long-term ecological understanding? Trends in Ecology & Evolution 19:39–45.

Holt, R. D., 1983a. Immigration and the dynamics of peripheral populations. Advances in Herpetology and Evolutionary Biology Pp. 680–694.

Holt, R. D., 1983b. Models for peripheral populations: the role of immigration. Pp. 25–32, in Population Biology. Springer.

Holt, R. D. and M. Barfield, 2011. Theoretical perspectives on the statics and dynamics of species borders in patchy environments. The American Naturalist 178:S6–S25.

Karlin, S. and R. Campbell, 1980. Selection-migration regimes characterized by a globally stable equilibrium. Genetics 94:1065–1084.

Lande, R., S. Engen, and B.-E. Sæther, 2009. An evolutionary maximum principle for density-dependent population dynamics in a fluctuating environment. Philosophical Transactions of the Royal Society of London B: Biological Sciences 364:1511–1518.

Lande, R., 2017. Evolution of stochastic demography with life history tradeoff’s in density-dependent age-structured populations. Proceedings of the National Academy of Sciences P. 201710679.

Lennon, J. J., J. R. Turner, and D. Connell, 1997. A metapopulation model of species boundaries. Oikos Pp. 486–502.

Lenormand, T., 2002. Gene flow and the limits to natural selection. Trends in Ecology & Evolution 17:183–189.

Levene, H., 1953. Genetic equilibrium when more than one ecological niche is available. The American Naturalist 87:331–333.

Levins, R., 1969. Some demographic and genetic consequences of environmental heterogeneity for biological control. American Entomologist 15:237–240.

Levins, R., 1970. Extinction. Some mathematical questions in biology.

Levins, R., 1979. Coexistence in a variable environment. The American Naturalist 114:765–783.

Long, Z. T., O. L. Petchey, and R. D. Holt, 2007. The effects of immigration and environmental variability on the persistence of an inferior competitor. Ecology letters 10:574–585.

Lorenz, E. N., 1963. Deterministic nonperiodic flow. Journal of the atmospheric sciences 20:130–141.

Luís, R., S. Elaydi, and H. Oliveira, 2011. Stability of a ricker-type competition model and the competitive exclusion principle. Journal of Biological Dynamics 5:636–660.

MacArthur, R. and E. Wilson, 1967. The theory of island biogeography. Princeton University Press.

May, R. M., 1972. Limit cycles in predator-prey communities. Science 177:900–902.

May, R. M., 1974. Biological populations with nonoverlapping generations: stable points, stable cycles, and chaos. Science 186:645–647.

May, R. M. and G. F. Oster, 1976. Bifurcations and dynamic complexity in simple ecological models. The American Naturalist 110:573–599.

Maynard Smith, J., 1970. Genetic polymorphism in a varied environment. The American Naturalist 104:487–490.

Pianka, E., 1970. On r-and k-selection. The American Naturalist 104:592–597.

Polechová, J., 2018. Is the sky the limit? on the expansion threshold of a species’ range. PLoS Biology 16:e2005372.

Rainey, P., A. Buckling, R. Kassen, and M. Travisano, 2000. The emergence and maintenance of diversity: insights from experimental bacterial populations. Trends in ecology & evolution 15:243–247.

Reddingius, J. and P. Den Boer, 1970. Simulation experiments illustrating stabilization of animal numbers by spreading of risk. Oecologia 5:240–284.

Ricker, W. E., 1954. Stock and recruitment. Journal of the Fisheries Board of Canada 11:559–623.

Roff, D., 1974. Spatial heterogeneity and the persistence of populations. Oecologia 15:245–258.

Roughgarden, J., 1971. Density-dependent natural selection. Ecology 52:453–468.

Ruxton, G. D., 1994. Low levels of immigration between chaotic populations can reduce system extinctions by inducing asynchronous regular cycles. Proc. R. Soc. Lond. B 256:189–193.

Sexton, J., P. McIntyre, A. Angert, and K. Rice, 2009. Evolution and ecology of species range limits. Annual Review of Ecology, Evolution, and Systematics 40:415–436.

Smith, H., 1998. Planar competitive and cooperative difference equations. Journal of Difference Equations and Applications 3:335–357.

Stone, L. and D. Hart, 1999. Effects of immigration on the dynamics of simple population models. Theoretical Population Biology 55:227–234.

Symonides, E., J. Silvertown, and V. Andreasen, 1986. Population cycles caused by overcompensating density-dependence in an annual plant. Oecologia 71:156–158.

Thomas, W. R., M. J. Pomerantz, and M. E. Gilpin, 1980. Chaos, asymmetric growth and group selection for dynamical stability. Ecology 61:1312–1320.

Tilman, D., R. M. May, C. L. Lehman, and M. A. Nowak, 1994. Habitat destruction and the extinction debt. Nature 371:65.

Turchin, P., L. Oksanen, P. Ekerholm, T. Oksanen, and H. Henttonen, 2000. Are lemmings prey or predators? Nature 405:562–565.

Turelli, M. and D. Petry, 1980. Density-dependent selection in a random environment: an evolutionary process that can maintain stable population dynamics. Proceedings of the National Academy of Sciences 77:7501–7505.

Wallace, B., 1975. Hard and soft selection revisited. Evolution Pp. 465–473.

Wright, S., 1931. Evolution in Mendelian populations. Genetics 16:97–159.

Yodzis, P., 2013. Competition for space and the structure of ecological communities, vol. 25. Springer Science & Business Media.

