## Supplementary text including all figures for "The influence of fluctuating population densities on evolutionary dynamics"

### Supporting Information: The influence of fluctuating population densities on evolutionary dynamics

**S4 Stronger selection weakens the impact of fluctuations** **16**

**S5 The effect of asymmetries – migration, selection, niche size** **17**

**S6 Robustness to logistic density dependence** **19**

**S7 Robustness to hard selection** **21**

**S8 Robustness to the order of the life cycle** **24**

**S9 Symmetric vs. unidirectional migration (density dependent dynamics)** **26**

**S10 Continent-island model with imposed fluctuations** **28**

### S1 Methods: Analytical analyses and resulting formulas

#### S1.1 Analyzing evolution in the absence of ecology

To reach a better understanding of the model's behavior we start by analyzing a simplification that takes into account evolution only. For this, we assume that population dynamics are in equilibrium and fix the population sizes to a constant value. We set

$$N_i(t) = c_i K. \tag{S1}$$

where $K$ denotes the carrying capacity of the entire habitat: $K = K_1 + K_2$, i.e. $c_i = K_i/K$. Thus, the formulas for the allele frequency after migration, $p_i'(t)$, equal

$$p_1'(t) = \frac{(1-m_{12})p_1(t)c_1 + m_{21}p_2(t)(1-c_1)}{(1-m_{12})c_1 + m_{21}(1-c_1)} \tag{S2}$$

because $K$ cancels out (and conversely for niche 2). I.e. we analyze a system of two recurrences for $p_1$ and $p_2$:

$$p_i(t+1) = p_i'(t) \cdot \frac{w_{iA}}{p_i'(t)w_{iA} + (1-p_i'(t))w_{ia}} \tag{S3}$$

We obtain the formula for a polymorphic equilibrium by solving a system of two equations for $p_1$ and $p_2$. Namely,

$$\begin{aligned} &\frac{(1-m_{12})p_1c_1 + m_{21}p_2(1-c_1)}{(1-m_{12})c_1 + m_{21}(1-c_1)} \cdot \\ &\cdot \frac{w_{1A}}{\frac{(1-m_{12})p_1c_1+m_{21}p_2(1-c_1)}{(1-m_{12})c_1+m_{21}(1-c_1)} w_{1A} + \left(1 - \frac{(1-m_{12})p_1c_1+m_{21}p_2(1-c_1)}{(1-m_{12})c_1+m_{21}(1-c_1)}\right) w_{1a}} + \\ &- p_1 \quad = \quad 0 \end{aligned}$$

and a second equation with 1 and 2 interchanged. Note, that we assume $m_{12}, m_{21} < \frac{1}{2}$.

We obtain the following equation for the allele frequency of $A$ at equilibrium, $p_i^*$:

$$
\begin{aligned}
p_1^* = 1 - \frac{1}{2c_1^2(1-m_{12})(1-m_{12}-m_{21})s_1^2 s_2} \cdot \Big( & m_{12}m_{21}s_1s_2 - m_{12}^2 m_{21}s_1s_2 + m_{12}m_{21}s_1^2 \\
& - m_{12}^2 m_{21} s_1^2 + s_1^2 s_2 - m_{21}s_1^2 s_2 - 2m_{12}s_1^2 s_2 + m_{12}^2 s_1^2 s_2 + m_{12}^2 m_{21} s_1^2 s_2 + \\
& + (1-c_1)^2(1-m_{12}-m_{21})s_1\big((1-m_{12})s_1s_2 - m_{21}(s_2(2-m_{12}(1-s_1)-s_1)-m_{12}s_1)\big) \\
& + (1-c_1)s_1\big(-2(1-m_{12})^2 s_1 s_2 + m_{21}(1-m_{12})(-2m_{12}s_1 + s_2(2-2m_{12}(1-s_1)+s_1)) \\
& + m_{21}^2(m_{12}s_1 - s_2(2-m_{12}-s_1+m_{12}s_1))\big) \\
& - c_1\sqrt{\begin{aligned}&(1-m_{12}-(1-c_1)(1-m_{12}-m_{21}))^2 s_1^2((1-m_{12})^2 s_1^2 s_2^2 + \\ &2m_{21}(1-m_{12})s_1s_2(m_{12}(2-s_2(1-s_1)-s_1)-s_1s_2) + \\ &m_{21}^2(s_1^2 s_2^2 - 2m_{12}s_1s_2(2-s_1-s_2+s_1s_2)+m_{12}^2(s_1+s_2-s_1s_2)^2))\end{aligned}}\,\Big)
\end{aligned}
\tag{S4}
$$

Note that the allele frequency of $a$ in niche 1 equals $1-p_1^*$ at equilibrium. Due to symmetry reasons, the equilibrium allele frequency of $a$ in niche 2 equals the above formula with all 1 and 2 interchanged. Thus, $p_2^*$ equals 1 minus the equation for $p_1^*$ with 1s and 2s switched.

In order to find conditions for stability we investigate the conditions for a protected polymorphism (i.e. the circumstances under which allele frequencies converge to fixation), as this approach simplifies the calculations and yields reasonably short equations: Instability of both monomorphic equilibria $p_1 = p_2 = 0$ and $p_1 = p_2 = 1$ implies stability of the polymorphic fixed point (because we consider a two-dimensional system of haploids: there is a unique, stable and globally attracting equilibrium (Karlin and Campbell, 1980)). So, we compute the Jacobi matrices $J_0$ and $J_1$ of the system at 0 and 1 respectively. Now, 0 is stable if and only if both eigenvalues of $J_0$ are in absolute value $< 1$. One can either compute the eigenvalues or use the following condition: Both eigenvalues of $J_0$ are in absolute value $< 1$ if and only if $|\mathrm{Tr}(J_0)| < 1 + \mathrm{Det}(J_0) < 2$. We did the latter for both $J_0$ and $J_1$ and obtained the equation that is stated in the article: for arbitrary migration rates $m_{12}$, $m_{21}$ and niche sizes $c_1$, $c_2$ the polymorphic equilibrium $(p_1^*, p_2^*)$ is stable if and only if

$$
s_1 > \frac{c_2 m_{21} s_2[1-m_{21}-c_1(1-m_{12}-m_{21})]}{[m_{21}+c_1(1-m_{12}-m_{21})]\cdot[c_1 m_{12} + c_2 s_2(1-m_{21})]} \tag{S5}
$$

and

$$
s_2 > \frac{c_1 m_{12} s_1[1-m_{12}-c_2(1-m_{12}-m_{21})]}{[m_{12}+c_2(1-m_{12}-m_{21})]\cdot[c_1 s_1(1-m_{12}) + c_2 m_{21}]} \tag{S6}
$$

For $m_{12} = m_{21} = m$ and $c_1 = c_2 = 1/2$ this simplifies to

$$
s_1 > \frac{m s_2}{m+(1-m)s_2} \quad \text{and} \quad s_2 > \frac{m s_1}{m+(1-m)s_1}. \tag{S7}
$$

Rearranging these inequalities yields $m/s_1 - m/s_2 < 1 - m$ and $-m/s_1 + m/s_2 < 1 - m$, which in turn leads to the condition stated in the main text.

Finally, we want to check whether the stability properties of the polymorphic equilibrium are the same in the absence of ecological dynamics and when they lead to stable population size – i.e. whether the above conditions represent an approximation or the exact solution. (By visual comparison such as in Fig. S1, we know that they are at least very similar.)

In order to do this, we compute the two eigenvalues of the system in the absence of ecology. (We choose not to include them in this text as they would take a few pages.) It turns out (computer algebra system) that the leading eigenvalue (which we determined visually by plotting both eigenvalues) of the simplified system is the same as the one eigenvalue of the complete system that captures the direct effects of selection and migration on evolution (we determined this eigenvalue visually, too – see Fig. S1). Therefore, the equations above are a precise condition for the stability of the polymorphic equilibrium in case of stable population dynamics, too.

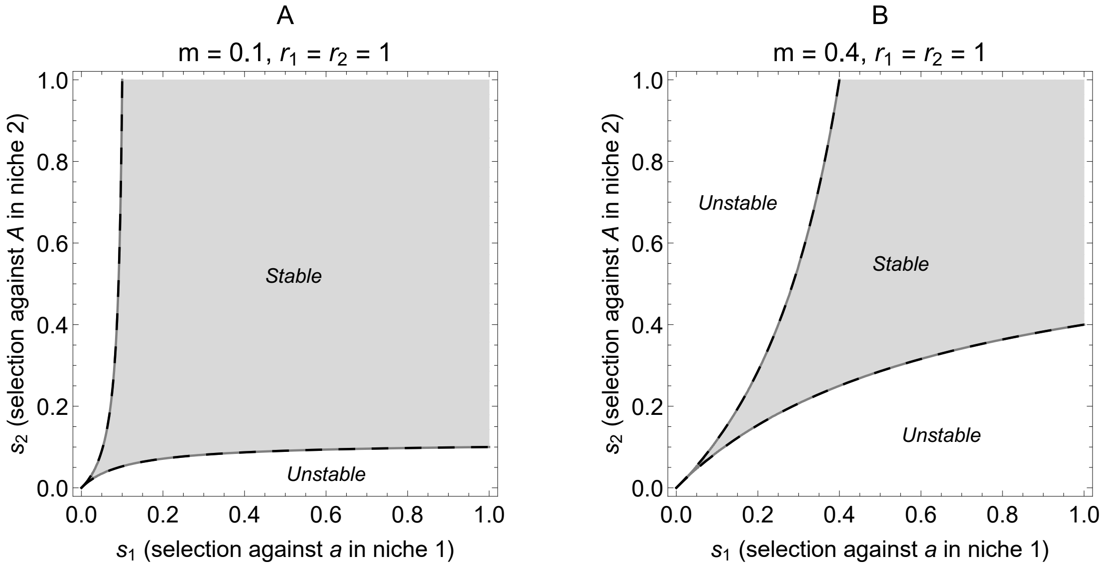

Fig. S1: **Stability of the polymorphic evolutionary equilibrium when ecological dynamics lead to stable population sizes.** For stable polymorphism to be maintained, the stronger the asymmetry in selection, the larger selection needs to be relative to migration (Maynard Smith, 1970; Bulmer, 1972). (A) weak migration; (B) strong migration. Migration is symmetric, $m = m_{12} = m_{21}$. Dark gray lines indicate instability due to monotonic divergence, i.e. where the eigenvalue, that describes the direct effect of selection and migration on the evolutionary equilibrium, is equal to 1. This eigenvalue coincides with the leading eigenvalue of the reduced system in the absence of ecological dynamics, which is depicted by the black dashed lines. The condition for stable coexistence is $m \cdot |1/s_1 - 1/s_2| < 1 - m$.

#### S1.2 Coupled system with density dependent population dynamics

According to the analysis of the evolutionary dynamics under stable population densities, we know the behavior of the system up to the first branching point of the ecological dynamics. We therefore want to reach a better understanding of that branching point. To asses its stability, we compute the Jacobian of the system. All results in this section are analytic, however, as the analysis yields extremely long and complex expressions, we choose to evaluate them and illustrate the results visually (see Fig. S4).

The Jacobian matrix has four eigenvalues, whereof two are of no interest as they are always smaller in absolute value than the other two. However, two of the eigenvalues are important and determine the stability of the equilibrium $(p_1^*, p_2^*, K_1, K_2)$. One of them, let's call it $\lambda_1$, captures the direct effects of selection and migration on the evolutionary equilibrium. It describes the stability of the polymorphic equilibrium as a function of $s_1$, $s_2$ and $m_{12}$, $m_{21}$ and is depicted in Fig. S1. The second eigenvalue of importance, $\lambda_2$, is shown in Fig. S4. It describes the ecological stability, that is to say, the influence of selection and migration on the first branching point of the population dynamics.

If selection and migration are symmetric, allele frequencies always converge to a polymorphic equilibrium. Thus, we can simply compute the Jacobian matrix of the system at the polymorphic equilibrium for $p_1$, $p_2$ and the population densities in equilibrium $N_1 = K_1$, $N_2 = K_2$. As polymorphism always exists, $|\lambda_1| < 1$ and we only need to consider $\lambda_2$ in order to understand the stability properties of the population dynamics. This is shown in Fig. S4A, B, C, D.

However, as the evolutionary dynamics influence the value of $r_i$ at the first branching point, we have to be a bit more careful as soon as we investigate asymmetric selection. Here, we need to distinguish the cases of convergence to fixation or convergence to polymorphism: In the regime where a stable polymorphism exists, we can use the eigenvalue $\lambda_2$ that we obtained in the previous step (by calculating the Jacobian at the polymorphic equilibrium). This regime is delimited by $|\lambda_1| < 1$. In the regime, where polymorphism doesn't exist, we need to compute $\lambda_2$ differently. We have to compute two other Jacobian matrices of the system: Once for the equilibrium $p_1 = p_2 = 0$ and once for the equilibrium $p_1 = p_2 = 1$. So, for parameter ranges that lead to fixation of $A$ we plot the $\lambda_2$ that we obtained from the Jacobian computed at $p_1 = p_2 = 1$ and vice versa for fixation of $a$. This is shown in Fig. S4E and F. The resulting picture in Fig. S4E and F is due to the change in $\overline{w}_i'(t)$ as a function of asymmetric selection. This can be seen easily in Fig. S5.

#### S1.3 Imposed fluctuations in the density of the focal niche

We consider a simplification of the model, where population densities are imposed: in niche 1, population size fluctuates between $K_1 + D_1$ and $K_1 - D_1$ ($K_1$, $D_1$ constant), while the density in niche 2 is fixed to $N_2 = K_1$ (see Fig. S2).

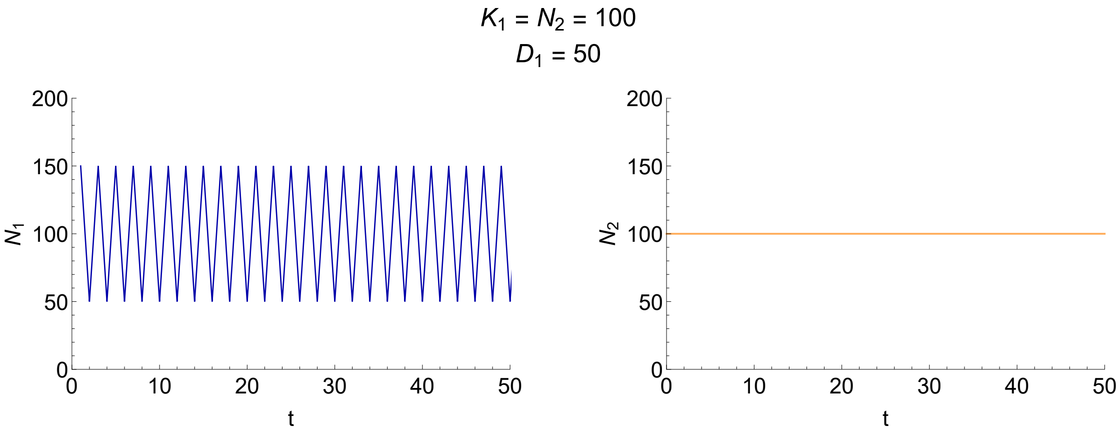

Fig. S2: **Imposing fluctuations $D_1$ in the population size of niche 1, while fixing the density in niche 2 to a constant value.** Population density in niche 1 periodically fluctuates between $K_1 + D_1$ and $K_1 - D_1$. Population size in niche 2 remains at $N_2$. (Parameters: $K_1 \pm D_1 = 100 \pm 50$, $N_2 = 100$.)

When migration is bidirectional, we are not able to predict the amount of variation maintained under fluctuating population densities, even when the fluctuations are imposed. Therefore, we assessed the stability analysis of the polymorphic equilibrium (Fig. 6 main text) again by investigating the protected polymorphism conditions instead (polymorphism is stable when both monomorphisms are unstable). As we trivially know the population densities $N_1 = K_1 \pm D_1$ and $N_2 = K_1$, we are able to analyze the stability of the monomorphic equilibria $p_1 = p_2 = 0$ and $p_1 = p_2 = 1$ (even in the presence of fluctuations).

In order to capture the fluctuations in population density, we analyze the stability of the monomorphisms under two consecutive generations. I.e. we compute the Jacobi matrices $J_0$ and $J_1$ of the iterated system (that describes the change in allele frequency after two generations) at 0 and 1. The leading eigenvalue of $J_0$, for example, reads

$$
\begin{aligned}
&\frac{1}{2(1+D/K(1-m))(1-D/K(1-m))(1-D/Km)(1+D/Km)(1-s_1)^2}\cdot\\
&\cdot\Big\{\Big(-1-(1-s_1)^2(1-s_2)^2\\
&\quad +m(6-8s_1+4s_1^2-8s_2+16s_1s_2-8s_1^2s_2+4s_2^2-8s_1s_2^2+4s_1^2s_2^2)\\
&\quad +m^2(DK^2-2(1+(1-s_1)(1-s_2)(2-3(s_1+s_2+s1s2))))\\
&\quad +m^3(-2(D/K)^2-2s_1+4s_1^2-2s_2+10s_1s_2-8s_1^2s_2+4s_2^2-8s_1s_2^2+4s_1^2s_2^2)\\
&\quad +m^4(DK^2-s_1^2-2s_1s_2+2s_1^2s_2-s_2^2+2s_1s_2^2-s_1^2s_2^2)\Big)\cdot(D/K)^2+\\
&+2-2s_1+s_1^2-2s_2+4s_1s_2-2s_1^2s_2+s_2^2-2s_1s_2^2+s_1^2s_2^2\\
&+m(-4+4s_1-2s_1^2+4s_2-8s_1s_2+4s_1^2s_2-2s_2^2+4s_1s_2^2-2s_1^2s_2^2)\\
&+m^2(4-4s_1+s_1^2-4s_2+6s_1s_2-2s_1^2s_2+s_2^2-2s_1s_2^2+s_1^2s_2^2)+\\
&+\sqrt{\begin{aligned}
&4(-1+(D/K)^2)(1-2m)^2(1+(D/K)^4(-1+m)^2m^2+\\
&+(D/K)^2(-1+2m-2m^2))(-1+s_1)^2(-1+s_2)^2+\\
&+\Big[2+(D/K)^4(-1+m)^2m^2-2s_1+s_1^2-2s_2+4s_1s_2-2s_1^2s_2+s_2^2-2s_1s_2^2+s_1^2s_2^2+\\
&\quad +m^2(-2+s_1+s_2-s_1s_2)^2-2m(2-2s_1(-1+s_2)^2+s_1^2(-1+s_2)^2-2s_2+s_2^2)+\\
&\quad -(D/K)^2\cdot\Big(2-2s_1(-1+s_2)^2+s_1^2(-1+s_2)^2-2s_2+s_2^2+\\
&\qquad -2m(3-4s_1(-1+s_2)^2+2s_1^2(-1+s_2)^2-4s_2+2s_2^2)+\\
&\qquad +2m^2(3+3s_1^2(-1+s_2)^2-5s_2+3s_2^2+s_1(-5+11s_2-6s_2^2))+\\
&\qquad +2m^3(-2s_1^2(-1+s_2)^2+s_2-2s_2^2+s_1(1-5s_2+4s_2^2))+\\
&\qquad +m^4(s_1+s_2-s_1s_2)^2\Big)\Big]^2
\end{aligned}}\ \Big\}
\end{aligned}
$$

The leading eigenvalue of $J_1$ is just as complicated. Therefore, these complex expressions are illustrated in Fig. 6 in the main text: in the parameter space where the leading eigenvalue of $J_1$ is in absolute value $< 1$, allele frequencies converge to fixation of the focal type $A$. Conversely, in the region where the absolute value of the leading eigenvalue of $J_0$ is smaller than 1 allele frequencies converge to fixation of the other type $a$. The region in between leads to polymorphism.

#### S2 Stability of the population dynamics

In the absence of evolution and migration, the ecological dynamics with Ricker's regulation are well understood. The population size converges towards a stable equilibrium as long as the intrinsic rate of increase is smaller than 2, which determines the first branching point in the absence of joint evolutionary dynamics (see Fig. S3A and S4A). As the growth rate increases above 2, the population size starts oscillating in a period-2 cycle due to overshooting of the carrying capacity and overcompensating density regulation (May and Oster, 1976). A further rise of the intrinsic rate of increase first leads to other period-doubling bifurcations and finally to chaotic behavior (see orange lines in Fig. S3A).

With migration and selection, the ecological dynamics change substantially. Fluctuations due to overcompensating density regulation in the focal niche are dampened by dispersal when the other niche maintains a stable density, as indicated by the blue lines in Fig. S3. Whereas the observation that migration between subpopulations may stabilize all kind of fluctuations has been known for some time (Den Boer, 1968; Reddingius and Den Boer, 1970; Roff, 1974), the extent of the dampening of potentially large and chaotic fluctuations is striking.

The stabilizing effect of constant migration into a Ricker regulated population has been explored in detail by Stone and Hart (1999): stabilization occurs because immigration pushes the population size away from zero. Chaotic behavior arises only for larger values of the intrinsic rate of increase, or not at all if migration is strong enough. Interestingly, even if the population size exhibits a phase of chaos, due to immigration it re-stabilizes for even higher intrinsic rate of increase, where fluctuations become periodic again (see Fig. S3, and Dey et al. 2014). Bidirectional exchange of individuals with a stable niche 2 leads to a slightly stronger stabilization than constant immigration explored by Stone and Hart (1999), because proportional emigration at high densities dampens the fluctuations (see section S9 for more detail).

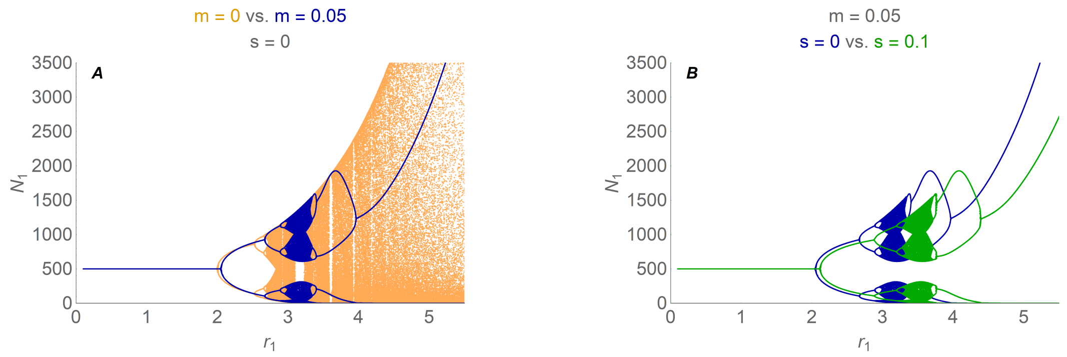

Fig. S3: **Migration and selection stabilize the population dynamics.** Bifurcation diagram showing the long-term behavior of the population size in niche 1 (while the population dynamics in niche 2 are stable with $r_2$ fixed to 1). (A) Migration from a stable environment dampens fluctuations and chaos which have arisen due to overcompensating density regulation. The standard Ricker model in the absence of migration and evolution is shown in orange. The blue lines depict $N_1(t)$ (after $t = 2^8 - 1$ to $2^8 + 1$ generations) as modeled in equation (4) with symmetric migration $m = 0.05$ (but no selection). (B) Selection shifts the branching points towards higher values: The green lines show the long-term behavior of the population dynamics with symmetric selection ($s_1 = s_2 = 0.1$). This stabilizing effect occurs because, here, the effective intrinsic rate of increase depends on the mean fitness which decreases with maladaptation. Note that the blue lines in (A) and (B) coincide. Parameters: $r_2 = 1$, $K_1 = K_2 = 500$.

In our model, selection also has a stabilizing effect on the population dynamics. This is because we assume throughout that the effective intrinsic rate of increase drops due to maladaptation. Immigration of maladapted individuals leads to a stronger stabilization in the population dynamics of the focal niche, because the more the population is maladapted the less it is able to exploit the potential rate of increase. As $\overline{w}'_i(t)$ decreases, the first branching point occurs for higher values of the intrinsic rate of increase only (see Fig. S3B).

In Fig. S4 we summarize the stability properties of the population dynamics as a function of the intrinsic rates of increase subject to selection and symmetric migration. In the case of symmetric selection (A, B, C, D) we simply observe the stabilizing effects of migration and selection as described above. However, asymmetry in selection (E, F) alters the behavior: The change in mean fitness $\overline{w}'_1(t)$ (which determines the value of $r_1$ at the branching point when migration is fixed) as a function of asymmetric selection is non-smooth and its properties depend on whether the equilibrium is polymorphic or not (see also Fig. S5).

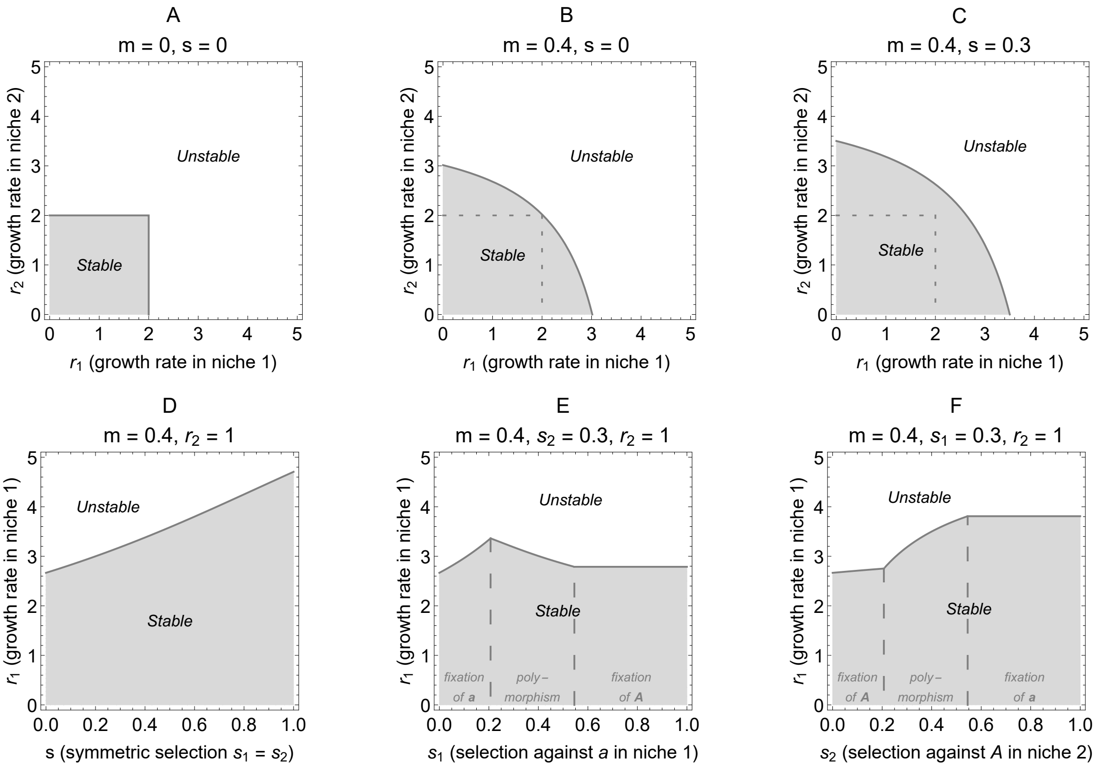

Fig. S4: **Stability of the ecological dynamics (first branching point).** Migration ($m$) and symmetric selection ($s$) stabilize the ecological dynamics. (A) In the absence of migration and selection, population size overshoots the carrying capacity for intrinsic rates of increase $r_1$, $r_2 > 2$ (for the Ricker model of population growth). (B) The stabilizing effect of migration on the population dynamics in niche 1 is stronger, the smaller the growth rate in the other niche, $r_2$, and vice versa. Strong migration between niches ($m = 0.4$) stabilizes the dynamics for $r_1$ up to 3 if $r_2$ is sufficiently small. However, if $r_1 = r_2$, migration has no stabilizing effect (the dashed lines indicate the stable region for $m = 0$, see A). (C, D) Here, strong symmetric selection decreases the mean fitness. This leads to a reduction in the effective intrinsic rate of increase and, thus, to a further stabilization of the population dynamics for large $r_1$, $r_2$. Note that in (A)—(D) the evolutionary equilibrium is always polymorphic due to symmetry in selection and migration. However, when selection is asymmetric, more complicated stability properties arise (E, F), as the asymmetry in allele frequencies at equilibrium influences the mean fitness after migration (see Fig. S5) and, thus, the value of $r_1$ at the first branching point: (E) Dashed lines indicate the existence of a polymorphic equilibrium. When $a$ is the only type present at equilibrium (left), increasing its local selective disadvantage $s_1$ corresponds to a decrease of the mean fitness in niche 1. In the middle region, a further increase of $s_1$ leads to a growing proportion of locally well adapted type $A$ individuals and mean fitness locally increases. When allele frequencies converge to fixation of type $A$ (right), mean fitness in the focal niche equals 1 and the branching point is determined by the stabilizing effect of migration only. (F) An increase in $s_2$, which determines the selection against $A$ in the other niche, leads to a drop of the equilibrium allele frequency of $A$ in both niches and, thus, to a decrease of the mean fitness in the focal niche (as the frequency of the locally maladapted type $a$ rises). The leading eigenvalue takes values between $-1$ and 1 within the gray areas, implying stability. The dark gray lines denote when it equals $-1$, which indicates instability due to oscillating divergence (see section S1.2 for explanation).

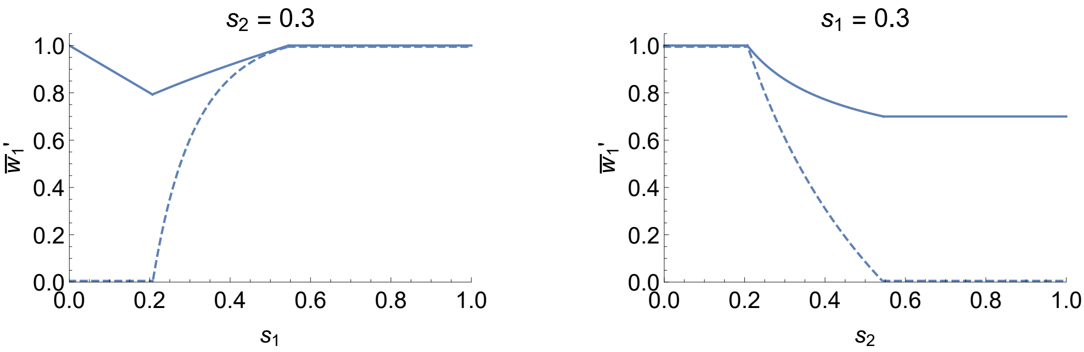

Fig. S5: **The mean fitness at evolutionary equilibrium changes as the asymmetry in selection is varied.** Solid lines indicate the values of the mean fitness after migration in the focal niche 1 (once evolutionary dynamics are in equilibrium), $\overline{w}_1'(t)$, as a function of one selection coefficient when the other is fixed. Dashed lines indicate the corresponding equilibrium allele frequency of $A$ in niche 1. The parameters here are the same as in Fig. S4E, F.

Left-hand side: Here, the selection against $A$ in the other niche, $s_2$, is fixed to 0.3, while the selective disadvantage of $a$ in the focal niche 1, $s_1$, is varied. For small values of $s_1$, $\overline{w}_1'(t)$ decreases, because no individuals of type $A$ are present at equilibrium: an increase of $s_1$ leads to a higher maladaption of $a$, which is the only type present. Then, a polymorphic equilibrium emerges and a further growth of $s_1$ leads to an increasing proportion of $A$, which are perfectly adapted to niche 1, and mean fitness increases. Finally, for large $s_1$, allele frequencies converge to fixation of $A$ and mean fitness equals 1, as $w_{1A} = 1$.

Right-hand side: Now, $s_1 = 0.3$ and we consider $\overline{w}_1'(t)$ as a function of the maladaptation of $A$ in niche 2, $s_2$. As long as $A$ is the only type present at equilibrium (for small $s_2$), mean fitness equals the fitness of $A$, which is 1 in the focal niche. As $s_2$ increases, the allele frequency of $A$ at equilibrium eventually decreases (in niche 2 and, thus, in niche 1, too). Mean fitness decreases as the locally maladapted type $a$ increases. For large values of $s_2$, allele frequencies converge to fixation of $a$ and, thus, mean fitness in niche 1 equals $1 - s_1 = 0.7$.

#### S3 Backward migration

The effect of fluctuations in population sizes is well approximated by changes in the mean backward migration.

Allele frequencies after migration equal

$$
\begin{aligned}
p_1'(t) &= \frac{(1-m_{12})N_1(t)}{(1-m_{12})N_1(t)+m_{21}N_2(t)} \cdot p_1(t) + \frac{m_{21}N_2(t)}{(1-m_{12})N_1(t)+m_{21}N_2(t)} \cdot p_2(t) \\
&= \frac{\alpha_1\, n(t)}{1+\alpha_1\, n(t)} \cdot p_1(t) + \frac{1}{1+\alpha_1\, n(t)} \cdot p_2(t) \\
p_2'(t) &= \frac{m_{12}N_1(t)}{(1-m_{21})N_2(t)+m_{12}N_1(t)} \cdot p_1(t) + \frac{(1-m_{21})N_2(t)}{(1-m_{21})N_2(t)+m_{12}N_1(t)} \cdot p_2(t) \\
&= \frac{\alpha_2\, n(t)}{1+\alpha_2\, n(t)} \cdot p_1(t) + \frac{1}{1+\alpha_2\, n(t)} \cdot p_2(t)
\end{aligned}
$$

where

$$
n(t) := \frac{N_1(t)}{N_2(t)}, \quad \alpha_1 := \frac{1-m_{12}}{m_{21}} \quad \text{and} \quad \alpha_2 := \frac{m_{12}}{1-m_{21}}
$$

Thus, when we define

$$
\mathcal{M}_1(t) = \frac{1}{1+\alpha_1\, n(t)} \qquad \textit{(“backward migration to deme 1”)} \tag{S8}
$$

$$
\mathcal{M}_2(t) = \frac{\alpha_2\, n(t)}{1+\alpha_2\, n(t)} \qquad \textit{(“backward migration to deme 2”)} \tag{S9}
$$

we obtain the allele frequencies after migration in terms of backward migration rates:

$$
p_1'(t) = \big(1-\mathcal{M}_1(t)\big) \cdot p_1(t) + \mathcal{M}_1(t) \cdot p_2(t) \tag{S10}
$$

$$
p_2'(t) = \mathcal{M}_2(t) \cdot p_1(t) + \big(1-\mathcal{M}_2(t)\big) \cdot p_2(t) \tag{S11}
$$

##### S3.1 Mean backward migration

When population densities fluctuate up and down with a period of 2, mean backward migration rates equal $\overline{\mathcal{M}}_i = 1/2 \cdot (\mathcal{M}_i(t-1) + \mathcal{M}_i(t))$. In the case of imposed fluctuations in niche 1, the formulas for the mean backward migration rates (over two generations) are known explicitly:

$$
\begin{aligned}
\overline{\mathcal{M}}_1 &:= \frac{1}{2} \cdot \left( \frac{1}{1+\alpha_1(1+D_1/K)} + \frac{1}{1+\alpha_1(1-D_1/K)} \right) \\
\overline{\mathcal{M}}_2 &:= \frac{1}{2} \cdot \left( \frac{\alpha_2(1+D_1/K)}{1+\alpha_2(1+D_1/K)} + \frac{\alpha_2(1-D_1/K)}{1+\alpha_2(1-D_1/K)} \right)
\end{aligned}
$$

$\overline{\mathcal{M}}_1$ and $\overline{\mathcal{M}}_2$ are plotted in the main text Fig. 4A as a function of $D_1/K$. The mean backward migration to the fluctuating niche, $\overline{\mathcal{M}}_1$, is an increasing, convex function of $D_1/K$, while the mean backward migration to the stable niche, $\overline{\mathcal{M}}_2$, is a decreasing, concave (yet close to linear) function of $D_1/K$. I.e. as imposed fluctuations in the focal niche increase, the mean backward migration to this niche grows fast, whereas the mean backward migration from it (to the other niche) declines slowly.

In order to understand the change in mean backward migration in the case of imposed fluctuations it is helpful to consider the following:

- $\mathcal{M}_1(n(t))$ and $1 - \mathcal{M}_2(n(t))$ are convex functions
- $1 - \mathcal{M}_1(n(t))$ and $\mathcal{M}_2(n(t))$ are concave functions

In the model with imposed fluctuations, we compare a scenario with constant $N_1(t) = K$ to a scenario with fluctuating $N_1(t) = K \pm D_1$, keeping the mean $\overline{N}_1 = K$ fixed. Since we also keep $N_2(t) = K$ constant, this means we let $n(t) = N_1(t)/N_2(t) = 1 \pm D_1/K$ fluctuate, keeping the mean $\overline{n} = 1$ constant. Then,

$$\big(\mathcal{M}_1(K + D) + \mathcal{M}_1(K - D)\big)/2 > \mathcal{M}_1(K)$$

and

$$\big(\mathcal{M}_2(K + D) + \mathcal{M}_2(K - D)\big)/2 < \mathcal{M}_2(K)$$

by Jensen's inequality. I.e. the averaged backward migration rate to the fluctuating niche, $\overline{\mathcal{M}}_1$, increases due to the fluctuations, while the averaged backward migration to the stable niche, $\overline{\mathcal{M}}_2$, decreases due to the fluctuations. This also explains that the niche with smaller fluctuations has an advantage (in terms of backward migration) – at least as long as we assume that the average population size remains constant. For the Ricker model, the fluctuations are not imposed, but a consequence of the overcompensation dynamics – and the mean population size is not constant.

#### S3.2 Residual effect of fluctuations in backward migration when approximated by its mean

In comparison to the approximation by a constant mean backward migration rate, fluctuations in backward migration rates lead to small "residual" changes in equilibrium frequency. This residual effect determines how well the complete system with fluctuating backward migration rates is approximated by their average. Here explain the approximation of fluctuating backward migration by its mean and show the this residual is small.

Why does the average backward migration approximate the impact of fluctuations in population densities on evolutionary dynamics so well? When population densities (resp. backward migration rates) fluctuate up and down, allele frequencies (exemplary in niche 1) after one generation equal

$$\begin{aligned} p_1'(t+1) &= (1-\mathcal{M}_1(t+1))\cdot p_1(t+1) + \mathcal{M}_1(t+1)\cdot p_2(t+1) \\ &= (1-\mathcal{M}_1(t+1))\cdot p_1'(t)\cdot w_{1A}/\overline{w}_1'(t) + \mathcal{M}_1(t+1)\cdot p_2'(t)\cdot w_{2A}/\overline{w}_2'(t) \end{aligned}$$

Close to the equilibrium, where $w_{iA}/\overline{w}_i'(t) \approx 1$, this approximates to

$$\begin{aligned} p_1'(t+1) &\approx (1-\mathcal{M}_1(t+1))\cdot p_1'(t) + \mathcal{M}_1(t+1)\cdot p_2'(t) \\ &= (1-\mathcal{M}_1(t+1))\cdot(1-\mathcal{M}_1(t))p_1(t) + (1-\mathcal{M}_1(t+1))\cdot\mathcal{M}_1(t)p_2(t)+ \\ &\quad + \mathcal{M}_1(t+1)\cdot\mathcal{M}_2(t)p_1(t) + \mathcal{M}_1(t+1)\cdot(1-\mathcal{M}_2(t))p_2(t) \\ &= \big(1-\mathcal{M}_1(t+1)-\mathcal{M}_1(t)+\mathcal{M}_1(t+1)\mathcal{M}_1(t)+\mathcal{M}_1(t+1)\mathcal{M}_2(t)\big)\cdot p_1(t)+ \\ &\quad + \big(\mathcal{M}_1(t+1)+\mathcal{M}_1(t)-\mathcal{M}_1(t+1)\mathcal{M}_1(t)-\mathcal{M}_1(t+1)\mathcal{M}_2(t)\big)\cdot p_2(t) \end{aligned}$$

As long as all the backward migration rates $\mathcal{M}_i \ll 1$, we can drop the higher order terms $\mathcal{M}_i(t)\mathcal{M}_i(t+1)$. Then, approximately

$$\begin{aligned} p_1'(t+1) &\approx \big(1-\mathcal{M}_1(t+1)-\mathcal{M}_1(t)\big)\cdot p_1(t) + \big(\mathcal{M}_1(t+1)+\mathcal{M}_1(t)\big)\cdot p_2(t) \\ &= p_1(t) + \big(p_2(t)-p_1(t)\big)\big(\mathcal{M}_1(t)+\mathcal{M}_1(t+1)\big) \end{aligned}$$

and thus mainly depends on the average backward migration rates.

When backward migration rates are approximated by their mean, the higher order terms $\mathcal{M}_i(t)\mathcal{M}_i(t+1)$ are omitted. Their impact is more significant when the fluctuations in backward migration rates $\mathcal{M}_i$ are strong, as this leads to really high values of either $\mathcal{M}_i(t)$ or $\mathcal{M}_i(t+1)$ (Fig. 4), and thus to an increase in the product $\mathcal{M}_i(t)\mathcal{M}_i(t+1)$.

Fig. S6 shows that the equilibrium allele frequencies decline under increasing fluctuations in backward migration rates – even if mean backward migration stays constant. However, this decline is very slight even when fluctuations in backward migration are strong. Therefore, the mean of the backward migration rates gives a robust approximation to the impact of fluctuating population sizes on evolutionary dynamics. The change in allele frequencies induced by fluctuations in population density is mainly due to a change in mean backward migration.

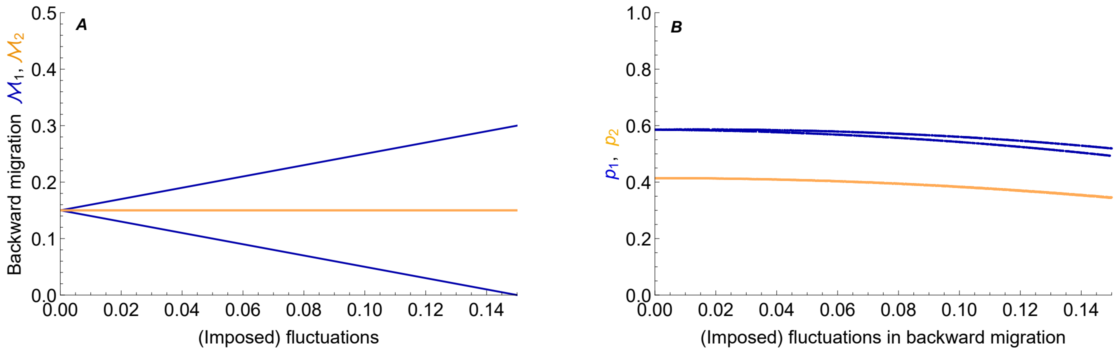

Fig. S6: **Residual effect of fluctuations in backward migration (A) on the equilibrium allele frequency (B).** The effect of fluctuations in population density on the evolutionary equilibrium is not completely explained by a change in the average backward migration rates. Even if the mean backward migration rate stays constant, equilibrium allele frequencies of the focal type decrease slightly when fluctuations in the backward migration to niche 1 increase (A, blue lines). (*c.f.* Fig. 3: the approximation of equilibrium frequencies deviates slightly when fluctuations in population density increase). Parameters: $s_1 = s_2 = 0.1$.

### S4 Stronger selection weakens the impact of fluctuations

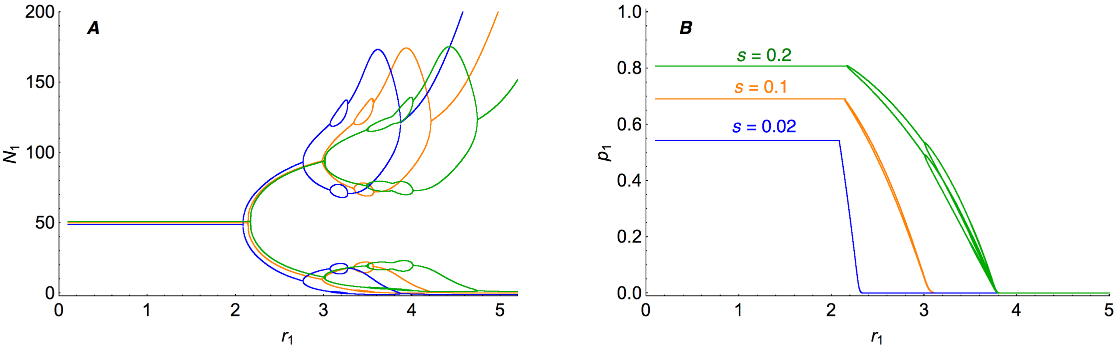

Fig. S7: **Symmetric selection counteracts the deleterious effect of fluctuations (B) and, together with migration, stabilizes the population dynamics (A).** This figure shows the dynamics at equilibrium as a function of the intrinsic rate of increase in niche 1 for three different values of selection ($s_1 = s_2 = 0.02$, 0.1, 0.2). (A) Stronger selection leads to a higher maladaptation on average; lower mean fitness then stabilizes the population dynamics in niche 1 because the effective intrinsic rate of increase decreases – the branching points shift to the right. As migration is slightly larger here comparing to Fig. 3, we observe a stronger stabilization of the fluctuations in the population size. (B) The extinction of the locally favorable type $A$ driven by large fluctuations in population size occurs for a higher $r_1$ under stronger selection. Mainly, this is because as selection increases, the timescale of evolution becomes comparable to the timescale of ecology, which decreases the impact of ecology on evolution. Note that the multiple branches in the curve for $s = 0.2$ are due to fluctuations in the allele frequency (not to bistability). Parameters: $m = 0.06$, $r_2 = 1$, $c_1 = 1/2$, $K = 100$.

### S5 The effect of asymmetries – migration, selection, niche size

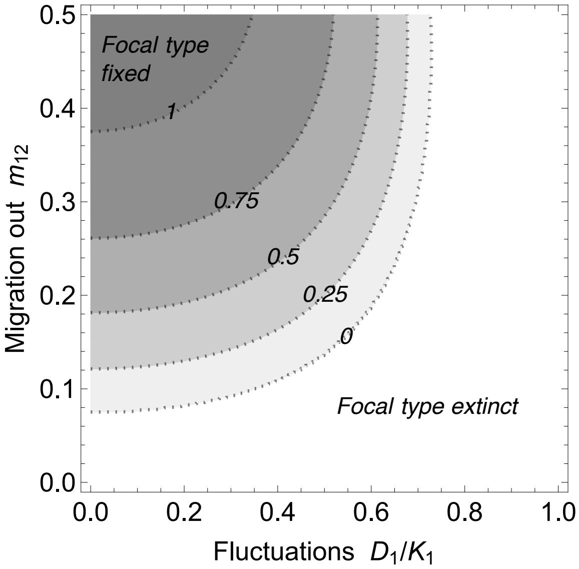

Fig. S8: **As fluctuations grow, more migration out of the niche (and then back in) may be essential for survival of the type adapted to the fluctuating niche**. Contours give the allele frequency in the focal niche, $p_1$: the focal type goes to fixation under high emigration ($m_{12} \to 0.5$) and low fluctuations ($D_1/K_1 \to 0$), whereas it goes extinct as fluctuations grow and emigration decreases. The other niche acts as a reservoir of the focal type, although the type is locally maladapted there. Hence, when immigration is constant (here, $m_{21} = 0.2$), more migration out always leads to an increase in frequency of the focal type. Parameters: $s_1 = s_2 = 0.1$, 10000 generations.

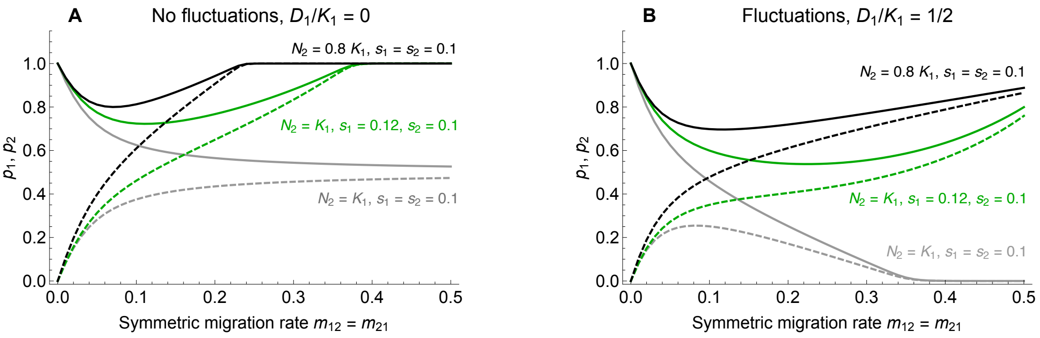

Fig. S9: **Increasing symmetric migration can be beneficial to the focal allele.** (A) Both niches have a stable population density. (B) The population density in the focal niche fluctuates with an amplitude of $D_1 = 1/2\,K_1$. (A, B) At first, as migration increases, the equilibrium frequency of the locally adapted allele ($p_1$) decreases in the focal niche (solid lines). Dashed lines give the equilibrium allele frequency of the focal type in niche 2, $p_2$. Yet, with stronger migration, the conditions for polymorphism to be stable become more sensitive to symmetry in selection coefficients and niche sizes (see S1.1 and Fig. 6) – and one or the other allele fixes. Therefore, if the focal niche is larger (black) or the selection against the invader is stronger (green), the focal allele will tend to swamp the other type in both niches as migration increases. Then, polymorphism is highest for intermediate migration rates, and increasing migration further leads to increase in equilibrium frequency of the focal allele. This recovery becomes more gradual and thus applies to a wider range of migration rates if the focal niche fluctuates, whereas the other one is stable (A *vs.* B).

#### S6 Robustness to logistic density dependence

We want to explore the system when population dynamics are modeled by logistic growth rather than Ricker's regulation. The equations for $p_1$ and $p_2$ remain unchanged, but the dynamics of the population density become

$$N_i(t+1) = N_i'(t) + N_i'(t) \cdot r_i \left(1 - \frac{N_i'(t)}{c_i K}\right) \cdot \overline{w}_i'(t). \tag{S12}$$

For the actual computations (Fig. S10 and S11) we furthermore force $N_i(t)$ to be non-negative. I.e. whenever the equations would lead to negative population sizes, we set them to zero instead and use that value to calculate the density in the next generation.

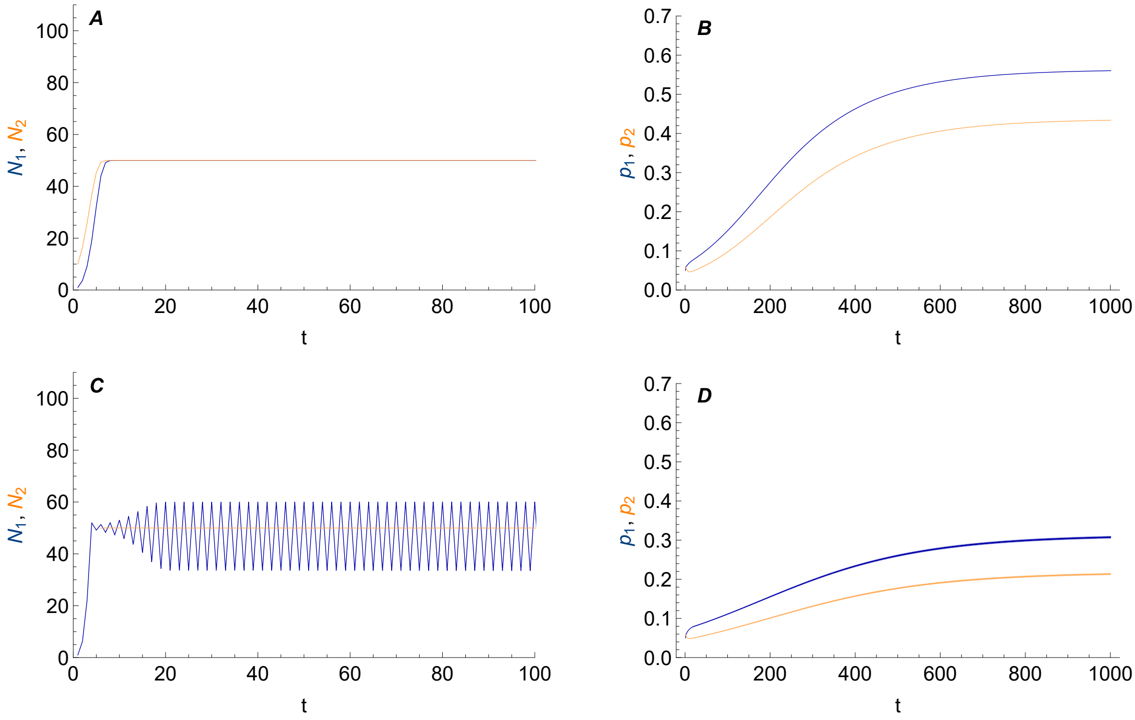

Fig. S10: **The qualitative behavior is preserved when Ricker's regulation is replaced by logistic growth.** The parameters here are the same as in Fig. 2 in the main text. Ecological dynamics modeled by logistic growth may affect the allele frequency at equilibrium. Compared to ecological dynamics that lead to stable population sizes (A, B), fluctuating population densities in niche 1 (C) lead to lower allele frequencies of A at equilibrium (D).

In Fig. S10 we see that the equilibrium allele frequency of the focal type $A$ is lower for fluctuating population dynamics (D) than it is for stable population dynamics (B). Hence, the decline in the equilibrium value of $p_1$ as a function of $r_1$ still exists when Ricker's regulation is replaced by logistic growth. Fig. S11 depicts the change in the allele frequency at equilibrium when $r_1$ is varied. The qualitative effect of population dynamics on evolution, i.e. the drop in allele frequencies of $A$, corresponds to what we observe in the main text, although there are quantitative differences. Comparing Fig. 3 (main text) and Fig. S11 (same parameters), we notice that with logistic growth the decrease in the equilibrium allele frequencies ($p_1$, $p_2$) with rising fluctuations is slower for weak fluctuations. This is because with logistic growth, the fluctuations in $N_1$ between first and second branching point are much weaker. Even in the chaotic regime, the fluctuations are weaker than under Ricker's regulation – hence equilibrium allele frequencies are still polymorphic when chaos arises in the ecological dynamics with logistic regulation.

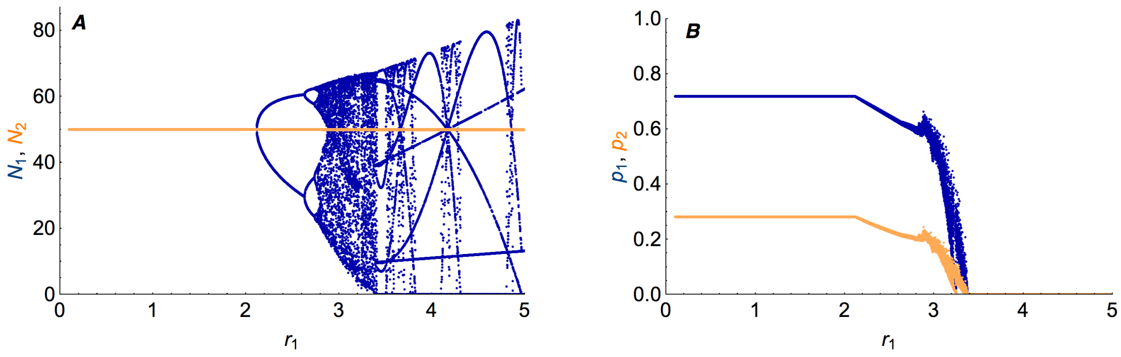

Fig. S11: **The qualitative long term behavior is preserved when Ricker's regulation is replaced by logistic growth.** The parameters here are the same as in the main text, Fig. 3 (only difference: here, generation time ranges from $2^{13}-1$ to $2^{13}+1$ as convergence is slower). (B) A shallow decrease in the allele frequencies of the focal type A ($p_1$ in niche 1, $p_2$ in niche 2) as small fluctuations arise is followed by are more drastic and rather chaotic one (as chaos arising in the population dynamics is transmitted to the evolutionary dynamics).

#### S7 Robustness to hard selection

The interactions between evolution and ecology can be modeled in many different ways. So far, we have assumed "soft selection" where carrying capacities are independent of the genetic composition of the population but the effective intrinsic rate of increase declines with maladaptation. Here, we will propose an alternative model with "hard selection" where mean fitness has an impact on both the effective intrinsic rate of increase and the carrying capacity.

We show that our main findings – namely, the change in equilibrium frequencies due to fluctuations in population densities – are robust to this alternative formalization.

Here, we assume that the growth rate of the population is given by the product of the population density regulation and the effects of genetic composition on its mean fitness $\overline{w}_i'(t)$. Thus, the population size in the next generation equals

$$N_i(t+1) = \overline{w}_i'(t) \cdot N_i'(t) e^{r_i\left(1-\frac{N_i'(t)}{K_i}\right)} \tag{S13}$$

In order to relate this formalisation to our soft selection model, we can rewrite this to the following:

$$N_i(t+1) = N_i'(t) \cdot e^{\tilde{r}_i(t)\left(1-\frac{N_i'(t)}{\tilde{K}_i(t)}\right)} \tag{S14}$$

where

$$\tilde{r}_i(t) = r_i + \log \overline{w}_i'(t) \quad \text{and} \quad \tilde{K}_i(t) = \frac{K_i}{r_i} \cdot \tilde{r}_i(t) = K_i \left(1 + \frac{\log \overline{w}_i'(t)}{r_i}\right)$$

Now we can see how the effective carrying capacity declines as maladaptation grows: $\tilde{K}_i(t)$ is an increasing function of $\overline{w}_i'(t)$ – the better is a population adapted to its niche, the higher is its effective carrying capacity. Only a perfectly adapted population reaches the potential carrying capacity $K_i$ ($\overline{w}_i'(t) \in (0,1]$ implies that $\log \overline{w}_i'(t)$ is negative).

The effective carrying capacity $\tilde{K}_i(t)$ depends also on the potential intrinsic rate of increase $r_i$. I.e. the mean fitness and the intrinsic rate of increase influence not only each other (as they do in our main model), but also the carrying capacity. Because there are more interactions between the parameters, the complexity of the system increases: a change in $r_i$ influences $\tilde{r}_i(t)$ and $\tilde{K}_i(t)$ directly. These, however, have an impact on the mean fitness $\overline{w}_i'(t)$, which again alters both $\tilde{r}_i(t)$ and $\tilde{K}_i(t)$, which again influence $\overline{w}_i'(t)$, and so forth.

The decrease in equilibrium allele frequency driven by fluctuations in population density is robust to this alternative formalization (Fig. S12). The deviations to our main model with soft selection can be readily understood – they follow from the increased complexity of this model with hard selection, where carrying capacity depends on intrinsic rate of increase and mean fitness. These effects are discussed in detail in the following paragraphs.

In this alternative model, the focal niche has a low effective carrying capacity $\tilde{K}_1(t)$ when the intrinsic rate of increase $r_1$ is small ($\log \overline{w}_1'(t)/r_1$ increases in absolute value). Then, the population size in the focal niche is small relative to the other habitat – and the focal type gets swamped by the migration from the larger niche 2. Therefore, the focal type converges to low frequencies when $r_1$ is close to zero in the alternative model.

However, both formalizations yield similar evolutionary results when the potential intrinsic rate of increase $r_1$ is not close to zero. For intermediate values of $r_1$, where population densities are not fluctuating, allele frequencies converge to a stable polymorphic equilibrium. However, in the alternative model with "hard selection" this polymorphism does not equal the one in the absence of ecology. In the alternative model, the allele frequency in the focal niche increases slightly with growing $r_1$ – because as long as there are no fluctuations in population density, the carrying capacity in niche 1 increases with growing $r_1$. Furthermore, as this leads to a higher proportion of the focal type, the mean fitness increases as well, which again augments the carrying capacity. Eventually, as $r_1$ increases, the effective carrying capacity of the focal niche can become larger than the carrying capacity in the other niche – until fluctuations arise.

In both models, the equilibrium frequency of the focal type decreases once fluctuations in the population density arise. These fluctuations occur for slightly smaller $r_1$ in the alternative model with hard selection, because the effective intrinsic rate of increase equals the sum of the (logarithmized) mean fitness and $r_1$ (as opposed to multiplication). Maladaptation has a weaker stabilizing effect on the population dynamics for large $r_1$ (as $r_1 \cdot \overline{w}_1'(t) \ll r_1 + \log \overline{w}_1'(t)$ when $r_1$ is big enough).

Under hard selection, the decrease in equilibrium frequencies with rising intrinsic rate of increase is more abrupt. Fluctuations in niche 1 decrease the local mean fitness, because they lead to a higher proportion of locally maladapted individuals – swamping increases. By decreasing the mean fitness, fluctuations drive a reduction in the effective carrying capacity of the focal niche. Thus, as maladaptation grows drastically with increasing fluctuations, the focal niche decreases in size relative to the other niche ($\tilde{K}_1(t) < \tilde{K}_2(t)$). In the presence of fluctuations, the focal type therefore converges to lower frequencies than it does in the main model – the effect of swamping becomes even stronger.

The differences between the two models in terms of decrease in equilibrium frequency depend on the influence of the mean fitness on the effective carrying capacity – and thus on the proportion of $\log \overline{w}_1'(t)/r_1$. On the one hand, when $r_1$ grows, the impact of the mean fitness on the carrying capacity declines (with $1/r_1$). However, under fluctuating population sizes, an increase in $r_1$ simultaneously leads to a drastic decrease in $\log \overline{w}_1'(t)$. All in all, this effect prevails and the focal allele converges to lower frequencies in the alternative model.

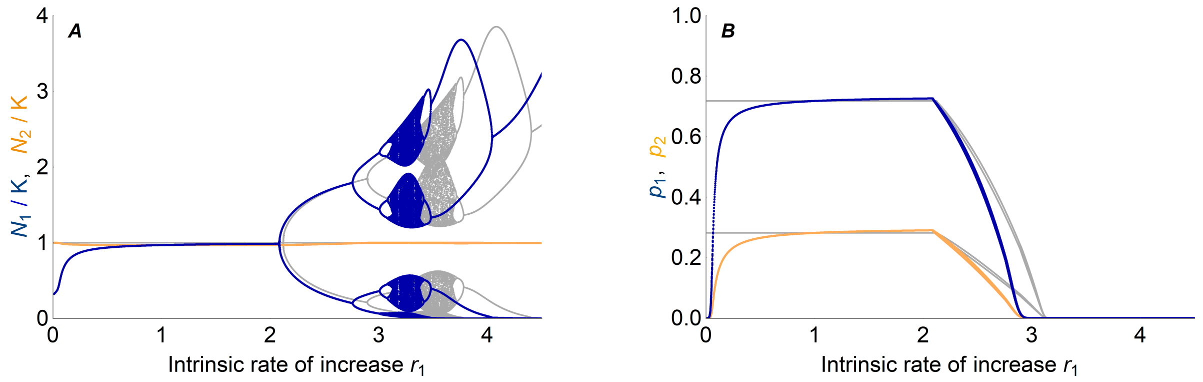

Fig. S12: **The decrease in equilibrium allele frequency induced by fluctuations in population density is robust to alternative formalizations – but with hard selection, when the carrying capacity depends on intrinsic rate of increase and mean fitness, additional effects emerge.** Here, we assume that the effective carrying capacity is proportional to $1 + \log \overline{w}_1'(t)/r_i$ and the effective intrinsic rate of increase equals $r_i + \log \overline{w}_i'(t)$ (see Eq. S14), which has the following effects: (A) Since the carrying capacity decreases with declining intrinsic rate of increase $r_1$, the population density in niche 1 (blue lines) converges for small $r_1$ to lower values than in the model with constant carrying capacity (grey lines). For large $r_1$, however, maladaptation has a weaker stabilizing effect on the population dynamics as it modifies the effective intrinsic rate of increase by addition rather than by multiplication: $r_1 \cdot \overline{w}_1'(t) < r_1 + \log \overline{w}_1'(t)$ when $r_1$ is large. (B) When the intrinsic rate of increase is small (and therefore the local carrying capacity is low), the focal type gets swamped by the migration from the second, larger habitat. When the intrinsic rate of increase is larger, both formalizations yield equivalent evolutionary results: for intermediate values of $r_1$, allele frequencies converge to a stable polymorphic equilibrium. Yet, in the alternative model this equilibrium deviates slightly from the one under soft selection, because the carrying capacity is still increasing with $r_1$. Once fluctuations in the population density arise with growing intrinsic rate of increase $r_1$, hard selection introduces a stronger decline in frequency of the type adapted to the fluctuating niche than soft selection with constant carrying capacity does (blue and orange *vs.* grey): as fluctuations lead to a higher proportion of locally maladapted individuals, local mean fitness decreases, which in turn reduces the carrying capacity of the focal niche – and increases the swamping. Parameters: $s = 0.1$, $m = 0.05$.

#### S8 Robustness to the order of the life cycle

In the main text, migration precedes selection and population growth. When the order in the life cycle is switched, i.e. when migration happens last – after selection and population growth, the equations turn into

$$N_1(t+1) = (1-m_{12}) \cdot \hat{N}_1(t) + m_{21} \cdot \hat{N}_2(t) \tag{S15}$$

and

$$p_1(t+1) = \frac{1}{N_1(t+1)} \cdot \left[ (1-m_{12}) \cdot p_1(t) \frac{w_{1A}}{\overline{w}_1(t)} \cdot \hat{N}_1(t) + m_{21} \cdot p_2(t) \frac{w_{2A}}{\overline{w}_2(t)} \cdot \hat{N}_2(t) \right] \tag{S16}$$

where

$$\overline{w}_i(t) = p_i(t) \cdot w_{iA} + (1-p_i(t)) \cdot w_{ia} \tag{S17}$$

equals the mean fitness in niche $i$ before migration, and

$$\hat{N}_i(t) = N_i(t) e^{r_i \left(1 - \frac{N_i(t)}{c_i K}\right) \overline{w}_i(t)} \tag{S18}$$

is the number of individuals in niche $i$ after selection (before migration).

Whether migration precedes selection and density regulation (as modeled in the main text) or migration succeeds selection and density regulation has no qualitative influence on the evolutionary dynamics. If selection (and population growth) occurred first, all individuals would reproduce according to the parameters of their current niche and offspring would migrate afterwards. This would lead to an increased coupling of the population dynamics between the niches: fluctuations in one niche would be transferred more strongly to the second niche (see Fig. S13).

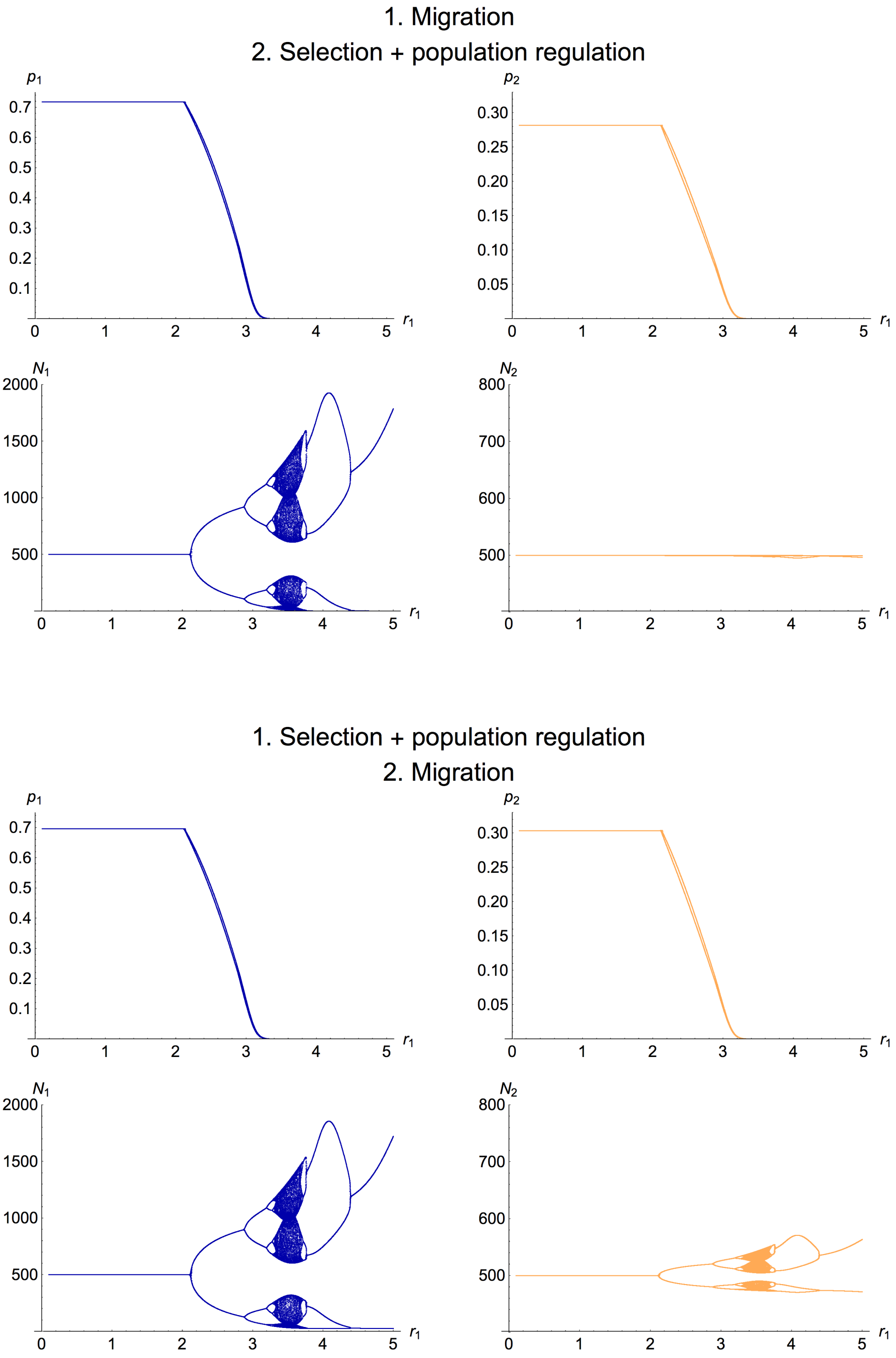

Fig. S13: **The influence of the order of the life cycle on the qualitative behavior of the system.** Upper figure: The system as it is modeled in the main text. Migration precedes selection and population regulation. Alternatively (lower figure), if migration comes last, we observe that the population size in niche 2 is coupled to the fluctuations in niche 1. How strongly the fluctuations are transmitted to the population density in niche 2 depends on the migration rate. Parameters: $r_2 = 1$, $s_1 = s_2 = 0.1$, $m_{12} = m_{21} = 0.05$.

#### S9 Symmetric vs. unidirectional migration (density dependent dynamics)

In the following, we explore how the coupled dynamics change when there is no migration out of niche 1 (i.e., $m_{12} = 0$, $m_{21} > 0$). Note that, as type $A$ individuals are locally maladapted in niche 2, they die out in this niche when it does not receive any immigrants. Therefore – and as the population density in niche 2 will be kept at carrying capacity ($r_2 = 1$) – the model then corresponds to a (potentially fluctuating) focal niche 1 receiving a constant monomorphic influx of locally maladapted individuals. Thus, this approach is akin to a continent-island model that includes population dynamics on the island.

**Effect of migration on population dynamics** Migration into a Ricker-regulated population has a stabilizing effect on the population dynamics (Stone and Hart, 1999), because it keeps the density away from zero. Yet, migration out ($m_{12} > 0$) can also stabilize population dynamics (see Fig. S14A), as the population size in the fluctuating niche 1 is pushed closer to its carrying capacity by symmetric migration rates ($m_{12} \approx m_{21}$ with niches of comparable size): When population size in the focal niche is high (fluctuating up), more individuals leave it than immigrants arrive from the other niche. Thus, there is a small net decrease in population density in niche 1 whenever the number of individuals is larger than the carrying capacity and the opposite is the case whenever population size is low (fluctuating down).

**Effects on evolutionary dynamics** The simplification of setting emigration from the focal niche to zero elucidates the role of the second niche as a refuge for the type favored in the niche undergoing strong fluctuations. With immigration only, the equilibrium frequency of the type better adapted to the fluctuating niche ($A$) is shifted to lower values comparing to bidirectional migration (see Fig. S14B). This is because when migration is unidirectional the allele frequency in niche 2 converges to fixation of type $a$: there is no immigration to niche 2 and $A$ is locally maladapted. Thus, immigration is monomorphic here, whereas it is polymorphic when migration goes both ways.

The second niche acts as a refuge for type $A$ even if selection is stronger than migration but only when the evolutionary dynamics are slow relative to the population dynamics (see Fig. S14B). When fluctuations are large (i.e. $r_1$ is large enough so that selection is weak relative to ecology), migration out of the niche is beneficial for the focal type as some proportion of the migrants survive in the neighboring habitat and may migrate back to the niche to which they are better adapted to, where they help to replenish the population in the growing phase. However, when population dynamics are very slow – for $r_1$ close to zero – the evolutionary dynamics becomes relatively strong. In this case, monomorphic immigration of the locally maladapted type $a$ leads to higher frequencies of the focal type $A$ than bidirectional migration does (Fig. S14B), because immigration and selection are both stronger than population regulation: As population size in niche

1 increases due to immigration when regulation is weak, the relative immigration decreases (since the absolute number of immigrants stays constant). This drop in relative immigration (which is monomorphic of the locally maladapted type) favors the locally well adapted type, because the timescale of selection remains unchanged.

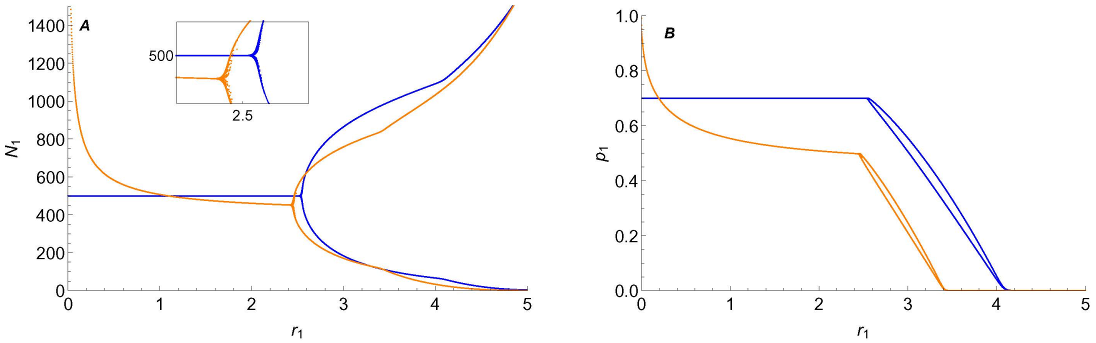

Fig. S14: **Unidirectional immigration has a weaker stabilizing effect on ecology and maintains less variation than a bidirectional exchange of individuals.** (A) The blue lines indicate the population dynamics in niche 1 with migration in both ways (with strong symmetric migration: $m = 0.2$). The outcome of population dynamics in niche 1 only with unidirectional immigration ($m_{21} = 0.2$) from the stable niche 2 ($r_2 = 1$) is shown in orange. The inset highlights the slightly stronger stabilization that occurs when migration is bidirectional. For $r_1$ small, immigration is stronger than density regulation, which leads to an overpopulation of niche 1. I.e. population density converges to a value above the carrying capacity whenever regulation is weak. In the extreme case of no regulation ($r_1 = 0$) population size in the focal niche goes to infinity over time, where $N_1(t+1) = N_1(t) + m_{21}N_2(t)$. However, for $r_1$ closer to the branching point, regulation is stronger and overcompensation occurs as a reaction to immigration which results in a slightly underpopulated niche 1. (B) The blue lines indicate the equilibrium allele frequency of $A$ in niche 1 with bidirectional migration, whereas the orange lines show the evolutionary outcome in niche 1 only with immigration from the stable niche 2 ($r_2 = 1$). The equilibrium allele frequency of $A$ is shifted to lower values when migration goes in one way only, as immigration is then monomorphic for type $a$. The exception is $r_1 \to 0$, where regulation is weak and, thus, outplayed by selection and immigration leading to an increase in equilibrium frequency of $A$. Parameters: $m_{12} = 0$ and $m_{21} = 0.2$ (orange) *vs.* $m_{12} = m_{21} = 0.2$ (blue), $s_1 = s_2 = 0.3$, $K = 1000$, $c_1 = 1/2$.

#### S10 Continent-island model with imposed fluctuations

We proceed by analyzing a continent-island model where we assume (imposed) fluctuations in the population size of the island. This simplification of the previously studied models yields simple, analytical results of the evolutionary dynamics on the island. Here, the formula for the allele frequency of the resident type equals

$$p_1(t+1) = p_1'(t) \cdot \frac{1}{p_1'(t) + [1 - p_1'(t)][1 - s_1]} \tag{S19}$$

where

$$p_1'(t) = \frac{p_1(t) N_1(t)}{N_1(t) + M_{21}} \tag{S20}$$

as immigration ($M_{21}$ is the total number of immigrants per generation) is monomorphic and, thus, $p_2 = 0$. The population size at the beginning of every life cycle equals $N_1(t) = K_1 + D_1$ or $N_1(t) = K_1 - D_1$, in subsequent generations. Here, we are able to solve for the equilibrium frequency of the resident type on the island. I.e. there are two relatively simple algebraic solutions to the equation

$$p_1(t+2) - p_1(t) = 0. \tag{S21}$$

Strictly speaking, as (equilibrium) allele frequencies fluctuate a bit due to the fluctuations in the population size, the frequency of type $A$ is converging to a two-cycle rather than a fixed point. Therefore, one of the two solutions to (S21) corresponds to the quasi-equilibrium frequency of $A$ in generations where population size on the island is high. The other, slightly smaller solution corresponds to the lower branch of the two-cycle. E.g. for the amount of variation maintained in generations where the population size on the island fluctuates up, we obtain the expression

$$\begin{aligned} p_1^* = {} & \frac{1}{2(K_1 - D_1)\Big((K_1 + D_1)(2 - s_1) + M_{21}(1 - s_1)\Big)s_1} \cdot \\ & \cdot \Bigg((K_1^2 - D_1^2)(2 - s_1)s_1 - M_{21}(2K_1 + M_{21})(1 - s_1)^2 + \\ & + \sqrt{((K_1^2 - D_1^2)(2 - s_1)s_1 - M_{21}(2K_1 + M_{21})(1 - s_1)^2)^2}\Bigg) . \end{aligned}$$

The conditions for persistence of the resident type $A$ (inequalities (9) and (10) in the main text) were obtained by setting both solutions for $p_1^*$ to zero. I.e. we calculated under which conditions type $A$ dies out (and then reversed the unequal signs).

The critical relative fluctuation size $(D_1/K_1)^*$ is the smallest fluctuation size that leads to extinction of the resident type.

$$\left(\frac{D_1}{K_1}\right)^* = \sqrt{\frac{1}{(2-s_1)s_1} - \left(\frac{1}{(2-s_1)s_1} - 1\right) \cdot \left(1 + \frac{M_{21}}{K_1}\right)^2}. \tag{S22}$$

If $(D_1/K_1)^*$ equals zero, or is a complex value, extinction of $A$ is certain – no matter if the population size on the island fluctuates or not.

The first order series of $(D_1/K_1)^*$ around $M_{21}/K_1 = 0$ is given by expression (S23). This is the rate at which the critical relative fluctuation size initially decreases as a function of the immigration rate $M_{21}/K_1$ (see Fig. S15): the approximation holds whenever $M_{21}/K_1$ is small. Approximation (S24) is true, if also $s_1$ is small.

$$\left(\frac{D_1}{K_1}\right)^* \approx 1 - \frac{M_{21}}{K_1}\left(\frac{1}{(2-s_1)s_1} - 1\right) \tag{S23}$$

$$\approx 1 - \frac{M_{21}}{K_1}\left(\frac{1}{2s_1} - 1\right) \tag{S24}$$

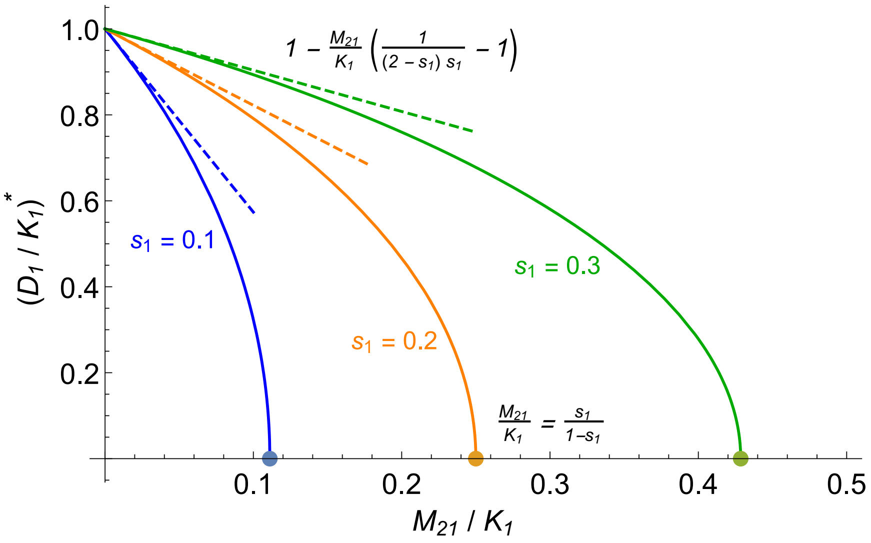

Fig. S15: **The decrease in critical relative fluctuation size as a function of the immigration rate is determined by the strength of selection against the immigrating type.** When immigration is weak (close to zero), the critical relative fluctuation size decreases with slope $-1/((2-s_1)s_1) + 1$. Hence, the weaker the selection $s_1$ against the immigrating type, the more detrimental proves its immigration $M_{21}/K_1$ to the resident type. At $M_{21}/K_1 = s_1/(1-s_1)$ the resident type dies out, no matter how strong are the fluctuations on the island.

### References

Bulmer, M., 1972. Multiple niche polymorphism. The American Naturalist 106:254–257.

Den Boer, P. J., 1968. Spreading of risk and stabilization of animal numbers. Acta biotheoretica 18:165–194.

Dey, S., B. Goswami, and A. Joshi, 2014. Effects of symmetric and asymmetric dispersal on the dynamics of heterogeneous metapopulations: two-patch systems revisited. Journal of theoretical biology 345:52–60.

Karlin, S. and R. Campbell, 1980. Selection-migration regimes characterized by a globally stable equilibrium. Genetics 94:1065–1084.

May, R. M. and G. F. Oster, 1976. Bifurcations and dynamic complexity in simple ecological models. The American Naturalist 110:573–599.

Maynard Smith, J., 1970. Genetic polymorphism in a varied environment. The American Naturalist 104:487–490.

Reddingius, J. and P. Den Boer, 1970. Simulation experiments illustrating stabilization of animal numbers by spreading of risk. Oecologia 5:240–284.

Roff, D., 1974. Spatial heterogeneity and the persistence of populations. Oecologia 15:245–258.

Stone, L. and D. Hart, 1999. Effects of immigration on the dynamics of simple population models. Theoretical Population Biology 55:227–234.
